# Evolutionary and phylogenetic insights from a nuclear genome sequence of the extinct, giant ‘subfossil’ koala lemur *Megaladapis edwardsi*

**DOI:** 10.1101/2020.10.16.342907

**Authors:** Stephanie Marciniak, Mehreen R. Mughal, Laurie R. Godfrey, Richard J. Bankoff, Heritiana Randrianatoandro, Brooke E. Crowley, Christina M. Bergey, Kathleen M. Muldoon, Jeannot Randrianasy, Brigitte M. Raharivololona, Stephan C. Schuster, Ripan S. Malhi, Anne D. Yoder, Edward E. Louis, Logan Kistler, George Perry

## Abstract

No endemic Madagascar animal with body mass >10 kg survived a relatively recent wave of extinction on the island. From morphological and isotopic analyses of skeletal ‘subfossil’ remains we can reconstruct some of the biology and behavioral ecology of giant lemurs (primates; up to ~160 kg), elephant birds (up to ~860 kg), and other extraordinary Malagasy megafauna that survived well into the past millennium. Yet much about the evolutionary biology of these now extinct species remains unknown, along with persistent phylogenetic uncertainty in some cases. Thankfully, despite the challenges of DNA preservation in tropical and sub-tropical environments, technical advances have enabled the recovery of ancient DNA from some Malagasy subfossil specimens. Here we present a nuclear genome sequence (~2X coverage) for one of the largest extinct lemurs, the koala lemur *Megaladapis edwardsi* (~85kg). To support the testing of key phylogenetic and evolutionary hypotheses we also generated new high-coverage complete nuclear genomes for two extant lemur species, *Eulemur rufifrons* and *Lepilemur mustelinus*, and we aligned these sequences with previously published genomes for three other extant lemur species and 47 non-lemur vertebrates. Our phylogenetic results confirm that *Megaladapis* is most closely related to the extant Lemuridae (typified in our analysis by *E. rufifrons*) to the exclusion of *L. mustelinus*, which contradicts morphology-based phylogenies. Our evolutionary analyses identified significant convergent evolution between *M. edwardsi* and extant folivorous primates (colobine monkeys) and ungulate herbivores (horses) in genes encoding protein products that function in the biodegradation of plant toxins and nutrient absorption. These results suggest that koala lemurs were highly adapted to a leaf-based diet, which may also explain their convergent craniodental morphology with the small-bodied folivore *Lepilemur*.

## Introduction

Madagascar is exceptionally biodiverse today. Yet the island’s endemic diversity was even greater in the relatively recent past. Specifically, there is an extensive ‘subfossil’ record of now-extinct Malagasy fauna, with some of these species persisting until at least ~500 years BP (before present)^1^. The late Holocene extinction pattern in Madagascar resembles other ‘megafaunal extinction’ patterns in that it is strikingly body mass-structured, with the majority of extinct subfossil taxa substantially larger than their surviving counterparts. For example, the average adult body mass of the largest of the ~100 extant lemur (primates) species is 6.8 kg^2^, well below that of the 17 described extinct subfossil lemur taxa, for which estimated adult body masses ranged from ~11 kg to an incredible ~160 kg^3^.

Despite a tropical and subtropical environment in which nucleotide strands rapidly degrade, in a select subset of Malagasy subfossil samples ancient DNA (aDNA) is sufficiently preserved for paleogenomic analysis^4–7^. In our group’s previous study^6^ we reconstructed complete or near-complete mitochondrial genomes from five subfossil lemur species, with population-level data in two cases (n=21 and n=3 individuals). As part of that work, we identified one *Megaladapis edwardsi* (body mass ~85 kg)^3,8^ sample UA 5180 (a mandible from Beloha Anavoha, extreme southern Madagascar; 1475 ± 65 cal yr. BP)^1^ with an especially high proportion of endogenous aDNA. We have subsequently performed additional rounds of extraction and sequencing of UA 5180 to amass sufficient data for studying the *M. edwardsi* nuclear genome.

We had two major analytical goals in the present study. First, to help reconstruct subfossil lemur behavioral ecology and evolutionary biology. Specifically, we can search unbiasedly across the genome for *Megaladapis-specific* signatures of positive selection at the individual gene level, and we can also search for striking patterns of genomic convergence with a set of biologically diverse extant mammals across sets of functionally-annotated genes. The results from these analyses may serve to extend current hypotheses or to offer potentially unexpected new insights into the evolutionary biology of *Megaladapis*.

Second, we aimed to resolve lingering uncertainty over *Megaladapis* phylogenetic relationships with other lemurs. At one point, a sister taxon relationship between *Megaladapis* and extant sportive lemurs (genus *Lepilemur*) was inferred based on craniodental similarities^3,9^. A different phylogeny was estimated, however, following the successful recovery of several hundred base pairs (bp) of the *Megaladapis* mitochondrial genome in several early aDNA studies^4,5^. Specifically, *Megaladapis* and the extant Lemuridae (genera *Eulemur, Lemur, Varecia, Prolemur*, and *Hapalemur*) formed a clade to the exclusion of *Lepilemur*. Our more recent aDNA study^6^ resolved a similar phylogeny, but with greater confidence (e.g. 87% bootstrap support) given the near-complete recovery of the *Megaladapis* mitochondrial genome (16,714 bp). Still, the mitochondrial genome is a single, non-recombining locus; in certain cases true species-level phylogenies are not reconstructed accurately from mitochondrial DNA only^10^. Most recently, Herrera and Dávalos (2016)^11^ estimated a ‘total evidence’ phylogeny by analyzing the combination of both morphological and genetic characters. Their result was dissimilar to each of the above phylogenies, instead supporting an early divergence of the *Megaladapis* lineage from all other non-*Daubentonia* (aye-aye) lemurs.

Because the nuclear genome is comprised of thousands of effectively independent markers of ancestry, we expected to achieve a more definitive phylogenetic result with our new *Megaladapis* paleogenome sequence. To distinguish among competing phylogenetic hypotheses, we also needed to generate new genome data for representatives of the extant Lemuridae and *Lepilemur* lineages, which we did for *Eulemur rufifrons* (red-fronted lemur) and *Lepilemur mustelinus* (greater sportive lemur), respectively. We aligned the three novel lemur genome sequences with those previously published for extant lemurs *Daubentonia madagascariensis* (aye-aye)^12^, *Microcebus murinus* (gray mouse lemur)^13^, and *Propithecus diadema* (diademed sifaka)^14^, and with 47 non-lemur outgroup species, for phylogenetic and evolutionary analyses.

## Results

We used a high-volume shotgun sequencing approach to reconstruct the *Megaladapis edwardsi* nuclear paleogenome. From the well-preserved *M. edwardsi* sample UA 5180 we had identified in our previous study^6^, we performed additional rounds of ancient DNA extraction (total extractions = 3), double-stranded library preparation (total libraries = 9), and massively parallel high-throughput ‘shotgun’ sequencing (total = 15 lanes on Illumina HiSeq 2000 and 2500 with 75 bp paired-end reads) to amass sufficient sequence data (total = 328 gigabases) for studying the nuclear genome despite the still relatively low endogenous nuclear DNA content (6.13%; see Methods; *SI Appendix*, **Figs. S1 and S2, Table S1**).

The size distribution^15,16^ and damage pattern^17–19^ (potentially damaged nucleotides were subsequently masked; see Methods) of putative *M. edwardsi* sequence reads were both characteristic of authentic ancient DNA (*SI Appendix*, **Figs. S3-S5**). Furthermore, we estimated a low 1.2% modern human DNA contamination rate among putative *M. edwardsi* reads (see Methods), consistent with or below that reported in other paleogenomic studies^20,21^ and overall contributing negligibly to the nuclear gene sequence reconstructions we report in this paper.

We aligned the set of quality-filtered and damage-masked *M. edwardsi* sequence reads to a version of the human reference genome (hg19) masked to contain only RefSeq gene exons ±100 bp. We used a conservative approach to reconstruct orthologous, single-copy *M. edwardsi* gene coding region sequences with 2X minimum sequence coverage per position (median proportion of sites reconstructed per gene = 0.29; *SI Appendix*, **Fig. S6, Table S2**). We also reconstructed exon sequences from shotgun sequence reads from the five modern lemur species (including two with data newly generated for this study) and a golden snub-nosed colobine monkey (*Rhinopithecus roxellana*) in our analysis^22^. All of these reconstructed sequences were integrated with a UCSC canonical gene exon alignment of 46 vertebrate species for an overall total alignment with 53 species.

### Reconstructing a nuclear genome-based phylogeny of extinct and extant lemurs

We used a genome-wide maximum likelihood approach^23^ to estimate the phylogenetic placement of *Megaladapis edwardsi* among primates. We first considered alignment data from n=896 genes for which at least 50% of *Megaladapis* sites were represented in our 2X minimum sequence coverage per position dataset (1.07 million bp in total) and estimated a single unrooted phylogeny from the concatenated alignment (Figure 1A). This extinct and extant lemur phylogeny, estimated from concatenated nuclear genome sequences, matches the previously-reconstructed mitochondrial genome-based phylogeny^6^.

**Figure 1.**
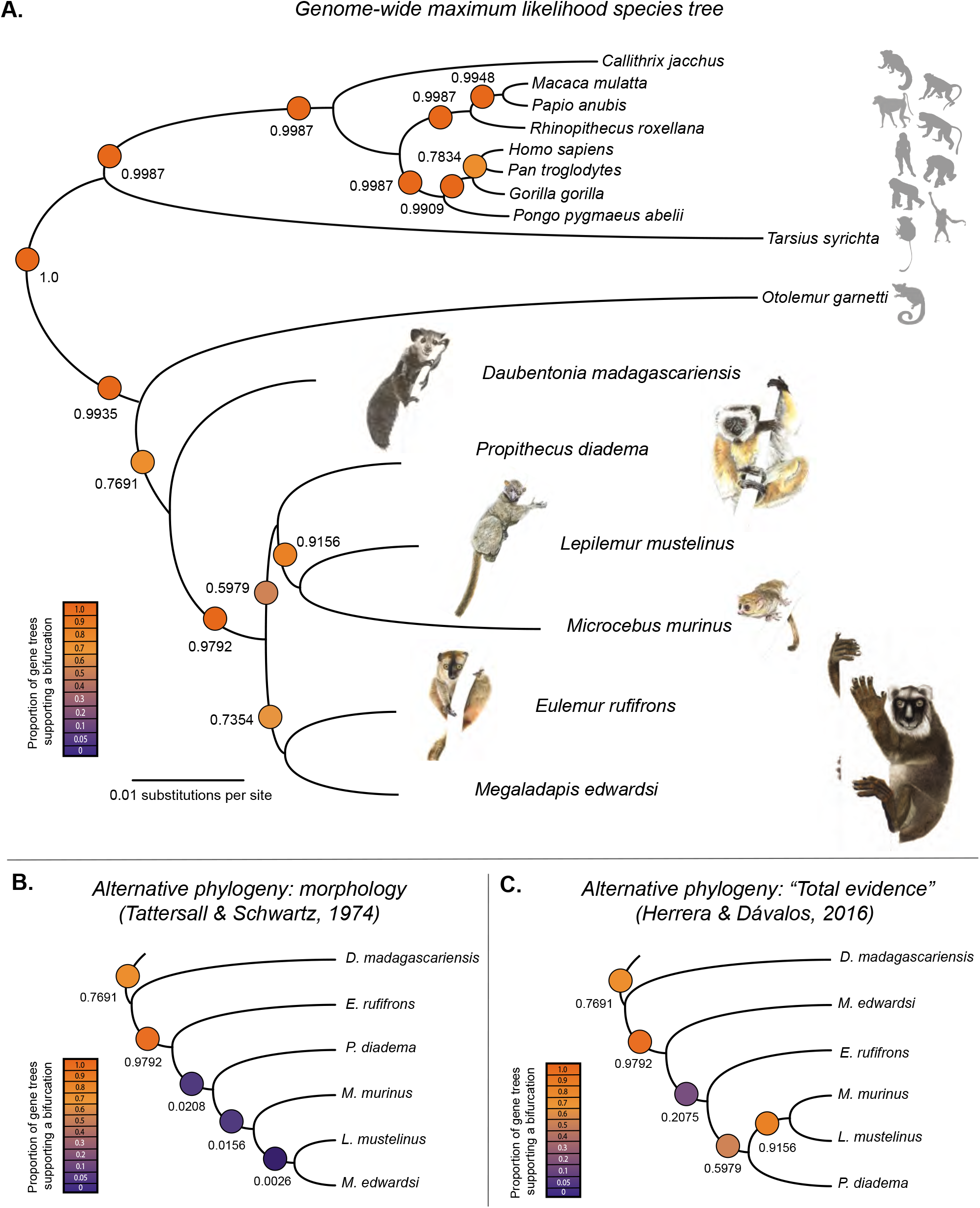
Phylogenetic analyses with the *Megaladapis edwardsi* nuclear genome sequence. A) Phylogeny estimated from maximum likelihood analysis of a concatenated alignment of the n=896 genes for which at least 50% of *M. edwardsi* sites were represented at minimum 2x sequence coverage (1.07 million bp in total). The heatmap and printed values represent the proportions of strong phylogenetic signal individual gene trees (a total of n=771 genes with ≥90% mean bootstrap support and ≥20% of sites present across all lemurs in the study) supporting each bifurcation. Watercolor illustrations by Joel Borgerson. Silhouette images courtesy of PhyloPic (see Methods for attribution details). B) The proportions of our strong phylogenetic signal individual gene trees that support each bifurcation in a previously hypothesized phylogeny inferred based on craniodental traits (Tattersall & Schwartz, 1974). C) The proportions of our strong phylogenetic signal individual gene trees that support each bifurcation in a previously published phylogeny based on the analysis of a combined morphological and mtDNA dataset (Herrera & Davalos, 2016).

Second, we analyzed a larger database of n=12,809 genes with aligned nucleotides present across at least 20% of the sites per gene across all lemurs in our study (including *M. edwardsi*). For each of these genes, we estimated an independent phylogeny using the same model as above and performed 100 bootstrap replicates. For each of these gene trees, the mean level of bootstrap support across all branch bipartitions was calculated as a measure of gene tree phylogenetic signal^24^. Among the 12,809 gene trees, the overall average mean bootstrap support value was 74.10% (s.d. = 12.65%; range = 7.69% to 98.85%; *SI Appendix*, **Fig. S7**).

We next considered the phylogenetic properties of the subset of individual gene trees with ≥90% mean bootstrap support. Of these n=771 “strong phylogenetic signal” individual gene trees, 191 (25%) exactly matched the full species tree based on the concatenated gene sequences. The species tree was well-supported at nearly every individual node (**Figure 1A**). The placement of *Megaladapis* as a sister taxon to *Eulemur* was supported in 567 out of the 771 strong phylogenetic signal gene trees (74%).

We also explicitly examined the level of support for the two alternative, previously reported phylogenies involving *Megaladapis*. First, scholars have hypothesized common ancestry for *Megaladapis* and *Lepilemur* to the exclusion of other lemurs based on a shared set of derived craniodental traits (e.g. the absence of permanent upper incisors, premolar proportional size similarity, and an expanded articular facet on the mandibular condyle) between these two taxa^9,25^. Yet a *Megaladapis*-*Lepilemur* sister taxon relationship was observed in only 2 of the 771 strong phylogenetic signal gene trees in our nuclear genome dataset (0.26%; **Figure 1B**).

Second, Herrera and Dávalos^11^ combined genetic data (for *M. edwardsi*: sequences from two mitochondrial genes^6^) and morphological trait variables (for *M. edwardsi*: n=169 traits) to reconstruct a “total evidence” phylogeny. Using their approach, *Megaladapis* was placed as a sister taxon to a clade of all other *non-Daubentonia* lemurs. This bipartition was observed in 160 of the 771 strong phylogenetic signal gene trees (20.75%; **Figure 1C**), the second-most observed result but a substantial ~3.5-fold reduction in support relative to the *Megaladapis-Eulemur* sister taxon relationship (**Figure 1A**).

### Evolutionary genomics

The *M. edwardsi* nuclear genome sequence contains a wealth of information about the evolutionary biology of this extinct species. Reliably equating between-species nucleotide differences to adaptive phenotypes is a considerable challenge regardless of genome quality^26,27^; and in our case here, the challenge is compounded by stochastic patterns of paleogenomic sequence coverage. Still, even with incomplete data, the vast expanse of the nuclear genome provides abundant opportunities to identify potential signatures of past natural selection. Combined with inferences of likely gene functions and pathways based on studies conducted in other species, these results can contribute to our understandings of *M. edwardsi* phenotypic form, function, and genetically-mediated behavior.

#### Nonsynonymous versus synonymous substitution rates

One comparative evolutionary genomics approach is to compare the ratios of the rates (*d*) of nonsynonymous (N; amino acid-changing) to synonymous (S; not amino acid-changing) substitutions (*d*_N_/*d*_S_) across a gene. While not all synonymous mutations are completely neutral with respect to function and fitness^28^, the fates of these mutations at least more closely reflect neutrality than those of nonsynonymous mutations. For the vast majority of genes in any interspecies comparison, *d*_N_/*d*_S_ ≪ 1, because the majority of nonsynonymous mutations are detrimental to fitness and are typically removed from populations by purifying selection. However, in rare cases, the repeated emergence of strongly adaptive nonsynonymous mutations at different positions along the same gene, leading to repeated fixation by positive selection, can lead to *d*_N_/*d*_S_ >> 1.

We used a maximum likelihood-based method implemented in the program PAML^29,30^ to estimate *d*_N_/*d*_S_ along each ancestral and terminal branch in our extant and extinct lemur genomic phylogeny. We restricted our analysis to the 3,342 genes with sufficient and high-quality sequence data for all lemurs and three outgroups (*H. sapiens, P. troglodytes*, and *G. gorilla*; see Methods). Because there are considerably fewer S than N sites per gene (e.g. in our dataset; 2.7 times fewer S sites overall), we further limited stochasticity in the *d*_N_/*d*_S_ statistic by computing a single, perlineage *d*_Sgenome_ value for use as the denominator in each gene-specific *d*_N_/*d*_S_ calculation^12^ (*d*_N_/*d*_Sgenome_) for that lineage.

We considered the 53 genes (1.6% of 3,342) with *Megaladapis* lineage-specific *d*_N_/*d*_Sgenome_ > 1.5 to be the strongest positive selection candidates for this extinct subfossil lemur in our dataset. When we tested whether this set of genes was significantly enriched for any known biological functions or biochemical pathways, we found none following multiple test correction^31^. Still, included among these 53 candidates were several individual loci with potentially intriguing links to hypotheses concerning *Megaladapis* evolutionary biology and behavioral ecology.

For example, *M. edwardsi* lineage *d*_N_/*d*_Sgenome_ = 2.83 for the growth hormone receptor (*GHR*) gene (N=7.8; S=2.0) whereas *d*_N_/*d*_Sgenome_ values for all other terminal and ancestral lemur branches range from 0.0 to 0.65 (**Figure 2A**; *SI Appendix*, **Fig. S8A**). Biological activity of growth hormone (GH) is mediated by interaction with the GHR protein. Genetic changes in the GH/GHR pathway can result in marked body size phenotypes^32–34^. Thus, the pattern of *Megaladapis-specific* positive selection in *GHR* marks this gene as a candidate contributor to the evolved gigantism in this lineage (estimated *M. edwardsi* body mass ~85 kg^3,8^ vs. maximum ~6.8 kg for any extant lemur^2^).

**Figure 2.**
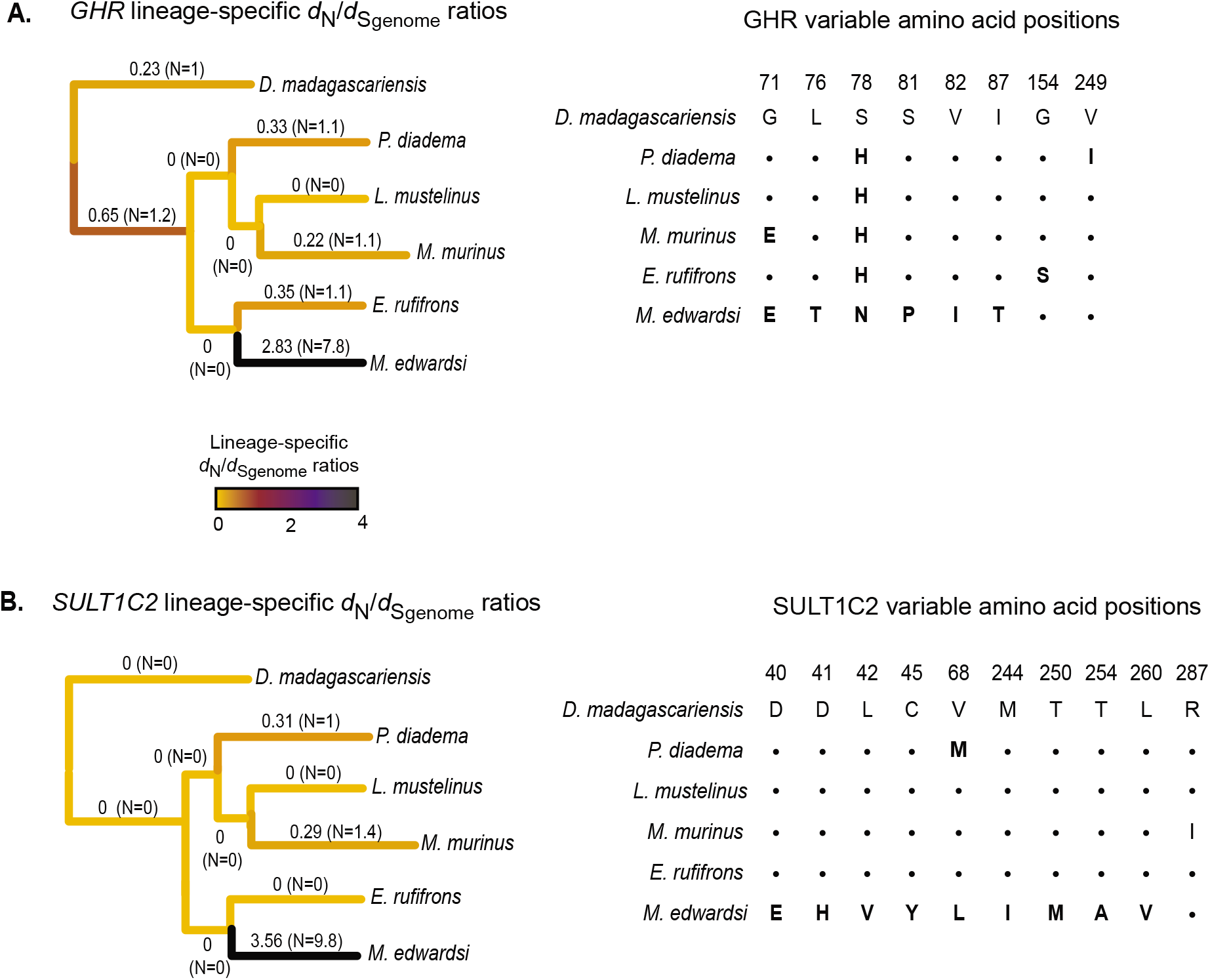
Lineage-specific *d*_N_/*d*_S_ ratios for *GHR* and *SULT1C2*. Using a maximum likelihood approach implemented in PAML, lineage-specific ratios of the rates (*d*) of nonsynonymous (N) vs. synonymous (S) substitution along ancestral and terminal branches estimated with a maximum likelihood-based approach for A) the growth hormone receptor (*GHR*) and B) sulfotransferase 1C2 (*SULT1C2*) genes. For each branch, the *d*_S_ denominator is based on the genome-wide synonymous substitution rate. *d*_N_/*d*_Sgenome_ estimates are recorded next to each branch and depicted by the heatmap. The estimated number of N substitutions for each branch are reported within the parentheses. Branch lengths shown are based on those from Figure 1A rather than these individual genes. For each gene, alignments of inferred amino acid residues for the encoded proteins are shown for all variable positions. Amino acid residues identical to those for *D. madagascariensis* are depicted with “.” and amino acid position numbers are based on the human reference sequence (hg19/GRCh37).

For the sulfotransferase 1C2 (*SULT1C2*) gene, *M. edwardsi* lineage *d*_N_/*d*_Sgenome_ = 3.56 (N=9.8; S=3.6) compared to a range of 0 to 0.35 for all other branches (**Figure 2B**; *SI Appendix*, **Fig. S8B**). SULT1C2 catalyzes reactions that detoxify xenobiotic compounds, including phenolics, to facilitate removal of potentially harmful metabolites from the body^35,36^. Phenolics are toxic compounds common in leafy plants^37^. Based on craniodental and postcranial gross morphology, biomechanical analyses, dental microwear and topographic analyses, and biogeochemistry, *M. edwardsi* is inferred to have been highly folivorous^38–41^. Thus, *SULT1C2* nonsynonymous substitutions may have been part of a suite of adaptations to folivory in the *Megaladapis* lineage (see more, below).

#### Convergent genomic evolution

The gene-by-gene *d*_N_/*d*_S_ approach presented above provides limited opportunity to identify signatures of past positive selection, as detection requires a history of repeated fixation of nonsynonymous substitutions within a gene beyond the background synonymous substitution accumulation rate. This combination can be especially rare on relatively longer branches such as the *Megaladapis* terminal lineage (i.e. estimated 27.3±4.2 MY divergence from last common ancestor with *Eulemur*^6^), resulting in likely high false-negative rates relative to the true occurrence of past positive selection.

Therefore, we also used a convergent evolution-based approach to identify potential signatures of positive selection on the *Megaladapis* lineage. We scanned across biological functional categories (i.e., groups of genes linked by known function based on the Gene Ontology (GO) database)^42^ to identify those functions with significantly higher proportions of convergent amino acid substitutions between *Megaladapis* and a distant species or clade relative to the genomewide rate of convergence. We performed this analysis using amino acid alignments of 21,520 genes for 53 total species, comprised of the six lemurs in our study (including *Megaladapis*) along with 47 non-lemur vertebrates (*SI Appendix*, **Fig. S9**). In combination with extensive available knowledge for many of the extant species in our dataset, these results can be used to develop or extend hypotheses of *Megaladapis* evolutionary biology and behavioral ecology.

Specifically, for each possible comparison between *Megaladapis* and a distant taxon (either an individual species or a clade of species), we searched for codon positions with the following pattern of convergent evolution: *Megaladapis* and the distant comparison taxon shared the same predicted amino acid, while the sister species to *Megaladapis (E. rufifrons*) and an outgroup lemur (*M. murinus*; we also performed separate analyses with *P. diadema; SI Appendix*, **Fig. S10**) shared a different amino acid, and the sister and outgroup species to the comparative taxon likewise shared a different amino acid. For each gene we also counted of the number of analyzable amino acid positions (see Methods). We then summed the numbers of convergent and analyzable sites across all genes represented in each GO term. For GO categories with ≥ 5 convergent amino acids we tested whether the proportion of convergent sites was significantly different than expected based on the genome-wide ratio.

Using this approach we performed 52 different comparisons between *Megaladapis* and a distant species/clade (*SI Appendix*). Per comparison, we identified an average of 0.54 (s.d.=0.90) GO categories significantly enriched for convergent amino acids at a low False Discovery Rate (FDR<0.05). Within any particular comparison, significant functional categories were often nested within other significant categories, as expected given the structure of the GO database.

Included among the most striking convergent evolution results were several patterns that may reflect *Megaladapis* adaptations to folivory. For example, between *M. edwardsi* and the golden snub-nosed monkey (*Rhinopithecus roxellana*), a colobine primate with a lichen- and leaf-specialized diet, there were 5 total convergent amino acid positions across 5 different hydrolase activity genes (GO:0016787; 8,535 total analyzable sites) versus an expectation of only 0.000057 convergent sites (genome wide convergent amino acids = 73; genome-wide analyzable positions = 1,273,496; Fisher’s exact test; P=0.00018; FDR=0.0054; **Figure 3A**). Among the identified hydrolase activity genes were *EXOG* and *ATP1A4*, which encode proteins involved in the metabolism of xenobiotics, which is critically important for many folivores given their exposure to plant secondary compounds^43^. Moreover, while families of genes involved in xenobiotic metabolism have expanded via gene duplication in golden snub-nosed monkeys^22^ and other herbivores^44,45^, in carnivores such genes are disproportionately pseudogenized^46^.

**Figure 3.**
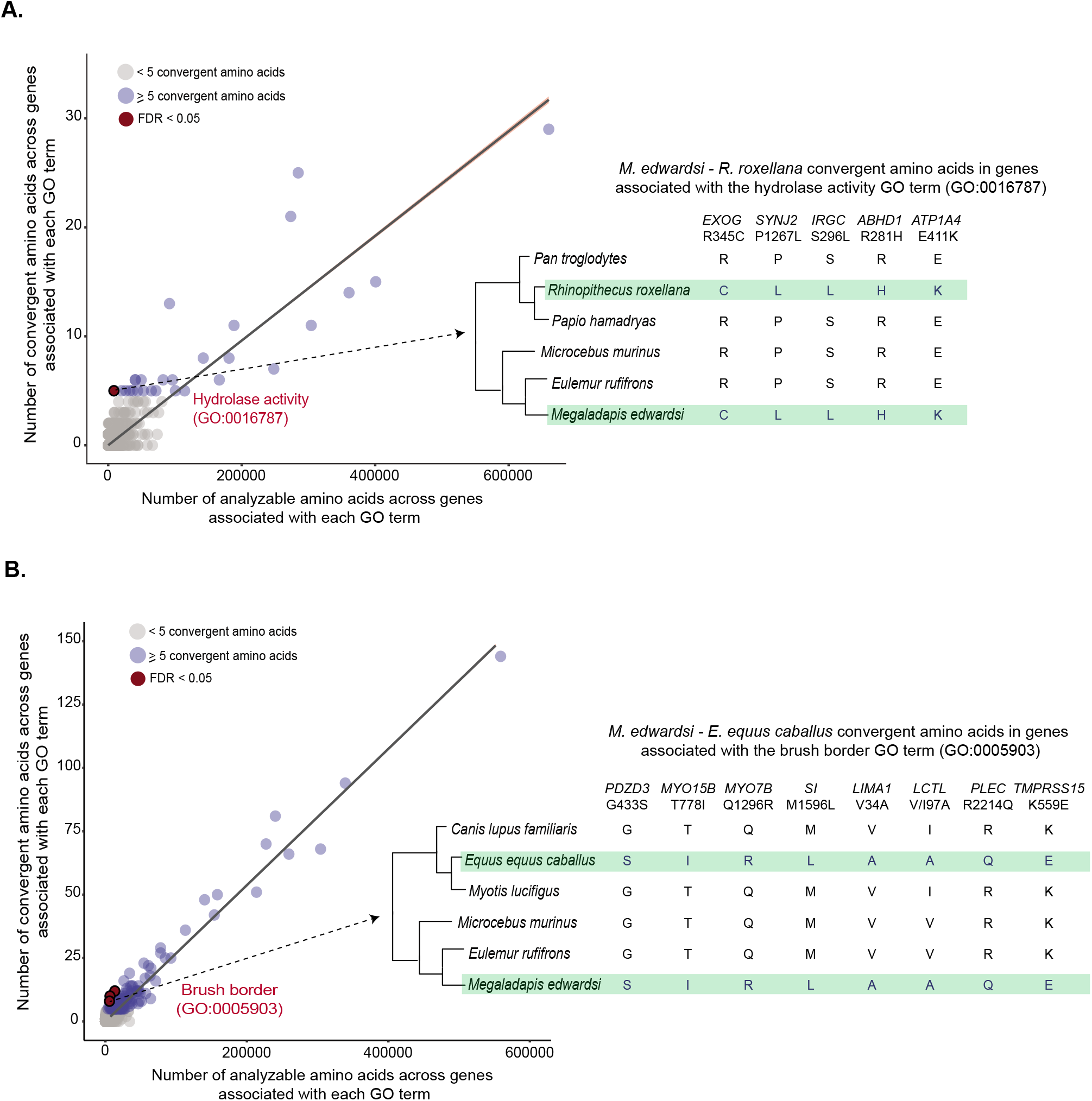
Convergent amino acid evolution between *Megaladapis edwardsi* and extant herbivores. Results from scans to identify Gene Ontology (GO) functional categories with unusual proportions (relative to genome-wide expectations) of inferred convergent amino acid positions between A) *M. edwardsi* and the folivore *R. roxellana* and B) *M. edwardsi* and the herbivore *E. equus caballus*. Convergent positions are those with identical residues between M. edwardsi and the comparison species, but for which the sister and an outgroup species (for each of the comparison species) share a distinct amino acid residue (shown at right). At left, the number of analyzable amino acid positions (aligned amino acids for all six species in the analysis plus identical residues in each sister species-outgroup pair) and convergent amino acid positions for each GO term. For terms with ≥ 5 convergent amino acids we tested whether the proportion of convergent sites was significantly different than expected based on the genome-wide ratio and computed false discovery rates (FDR) to account for the multiple tests. For two highlighted GO terms, all convergent amino acid positions between *M. edwardsi* and the comparison species along with gene name and position (based on the human reference sequence) are shown.

We also identified 8 total convergent amino acids across 8 different brush border genes (GO:0005903) between *Megaladapis* and horse (*Equus caballus*) versus an expectation of 0.0003 convergent sites (brush border gene analyzable sites = 5,787; genome-wide convergent positions = 316; genome-wide analyzable positions = 1,058,758; Fisher’s exact test; P=0.00046; FDR=0.0307; **Figure 3B**). The brush border is the microvilli-covered surface of epithelial cells, for example the intestinal lining, which helps to facilitate the absorption and hydrolysis of nutrients (via brush border enzymes embedded in the microvilli)^47^. Brush border genes with *Megaladapis*-horse convergent amino acids include *LIMA1*, which encodes an actin-binding protein with a role in cholesterol homeostasis^48^, and the microvilli myosin-encoding gene *MYO7B*, which maintains brush border action^49^. The digestive biology of these brush border proteins in horses is incompletely known, but the connection between the herbivorous diet of horses and the proposed specialized folivory of *Megaladapis* warrants further investigation into the functional impacts of these convergent amino acid changes^50–52^.

## Discussion

For this study we generated the first nuclear genome sequence dataset for an extinct non-hominin primate species. Paleogenomic approaches have immense potential for helping to resolve phylogenetic relationships and for insights into the evolutionary biology of now-extinct taxa and ancestral clades^53^. Our study follows the recent analysis of a nuclear genome sequence from a ~5,800 years BP baboon (extant *Papio ursinis*)^54^, the sequencing of a mitochondrial genome and five nuclear genes from an extinct Caribbean monkey (*Xenothrix mcgregori*)^55^, and prior mitochondrial DNA sequencing studies of multiple extinct subfossil lemur species^4–7^. In addition to paleogenomics, we are following continuing developments in the field of paleoproteomics^56^ for similar insights from samples with inadequate ancient DNA preservation, including those considerably older. An exciting recent paper presenting and analyzing the enamel proteome of the extinct orangutan relative *Gigantopithecus blacki* demonstrated this point^57^.

For the present study, we felt fortunate to generate *Megaladapis edwardsi* nuclear genome sequence data. Madagascar’s tropical and sub-tropical conditions severely challenge ancient DNA preservation. To date our ancient DNA laboratory has screened multiple hundreds of extinct subfossil lemur samples (many had been collected previously for non-ancient DNA analyses). Yet we have considered endogenous DNA preservation sufficient in only two samples to attempt (at least with current technology) shotgun sequencing of the nuclear genome. The *M. edwardsi* sample UA 5180 studied here was the best preserved.

### Phylogenetic resolution of a rapid lemur radiation with incomplete lineage sorting

Our ability to analyze sequence data from thousands of loci from across the *M. edwardsi* nuclear genome helped us to resolve ongoing extant-extinct lemur phylogenetic uncertainty, particularly the branching order of Lemuridae-Megaladapidae, Lepilemuridae-Cheriogalediae and Indriidae. Prior analyses of mtDNA sequence data from *M. edwardsi* and extant lemurs showed that *Megaladapis* and extant *Lepilemur* were likely not sister taxa^4,6^, as previously had been hypothesized based on morphological similarities^3,9^. Both of these phylogenetic reconstructions positioned *Megaladapis* distinctly from yet another, more recent phylogenetic analysis that was based on an extensive morphological plus mtDNA combined dataset^11^.

A sister taxon relationship between *Megaladapis* and extant Lemuridae (represented in our study by *Eulemur rufifrons*) was robustly supported in our nuclear genome-based analysis (**Figure 1A**) relative to alternative phylogenies (**Figure 1B-C**). This result is consistent with the prior phylogenetic reconstructions based on mtDNA sequences only. Our nuclear phylogeny does not support a close relationship of the Megaladapidae and the Lepilemuridae; instead, the latter is the sister to the Cheirogaleidae, and the lepilemurid-cheirogaleid clade is the sister to a clade comprising the Archaeolemuridae, Indriidae and Palaeopropithecidae.

We propose two non-mutually exclusive explanations for the past phylogenetic inference discrepancies. First, based on patterns of dental microwear^38,40,58^, dental topography^41^, craniodental features^9,25^, infraorbital foramen size^59^ and isotopic data^60–62^ with further support from our evolutionary genomic results (see below), *Megaladapis* was likely a specialized folivore. Meanwhile, the diets of sportive lemurs (*Lepilemur* spp.) are also highly folivorous^63–65^. *Megaladapis*-*Lepilemur* morphological similarities may thus represent convergent biological adaptations to similar behavioral ecology, rather than shared inheritance from a common ancestor. For example, the absence of upper incisors in both taxa could represent convergent adaptation to folivory in the context of a plesiomorphic lower toothcomb. Such processes could affect phylogenetic analyses based on morphological features.

Second, a rapid early diversification of lemur lineages (other than *Daubentonia*) occurred ~34 million years ago^6^. Potentially, this rapid radiation was triggered by the Eocene-Oligocene extinction event, a period of dramatic climate shift (global cooling) and flora/fauna turnover (forest reduction, niche fragmentation)^6,66,67^. Alternatively, based on recent African fossil evidence, there may have been two separate lemur colonizations of Madagascar - one, by an ancestor exclusive to the *Daubentonia* lineage and another by an ancestor of all non-*Daubentonia* lemurs (which could have occurred at ~34 MYA during the Cenozoic)^68^. Regardless, rapid radiations like this likely complicate lemur phylogenetic reconstructions.

Specifically, within a closely timed radiation, a proportion of ancestral genetic variants may remain polymorphic across multiple lineages through the duration of splitting events, to only subsequently become fixed – potentially with a fixation pattern that is not representative of species-level relationships. This “incomplete lineage sorting” process^69,70^ can lead to conflicting locus-to-locus phylogenetic signals, thereby resulting in a minority of gene trees differing from the overall species tree.

Incongruences due to incomplete lineage sorting are not uncommon among primates. For example, this phenomenon has been well-documented for humans, chimpanzees, and gorillas. Across the autosomal nuclear genome of these species, ~30% of alignments support incongruent branching orders of (chimpanzee, (human, gorilla)) or (human, (chimpanzee, gorilla) instead of the true species order (gorilla, (human, chimpanzee))^71^. Indeed, this finding is replicated by our own gene tree / species tree phylogenetic analysis, with results from 604 of 771 genes (78.3%) supporting the true (gorilla, (human, chimpanzee) phylogeny (Figure 1A) versus results from 78/771 genes (10.12%) supporting (chimpanzee, (human gorilla)) and 89/771 (11.54%) supporting (human, (gorilla, chimpanzee)) incongruent branching orders.

Using the same set of genes, we observed a similar signature of incomplete lineage sorting among lemur clades involving *Megaladapis*. Specifically, the ((*Megaladapis*, *Eulemur*), all other non-*Daubentonia* lemurs) typology was supported by 567 of the 771 gene trees with strong phylogenetic signal (73.5%; **Figure 1A**). The second-most common branching order involving *Megaladapis* (*Daubentonia*, (*Megaladapis*, all other lemurs)) was supported by 160/771 (20.8%) of gene trees (**Figure 1C**). This signature of incomplete lineage sorting strongly supports the notion of a rapid radiation among non-*Daubentonia* lemurs on Madagascar, possibly immediately following either a mass extinction event^6,72^ or a non-*Daubentonia* lemur colonization of the island^68^.

### Evolutionary genomic reconstruction of *Megaladapis* as a large-bodied specialized folivore

We approached our evolutionary genomic analyses with care, and we suggest cautious interpretation of the results. The *Megaladapis* lineage branch length is relatively long, making it difficult to identify individual genes with histories of positive selection based on the detection of excessive nonsynonymous substitution fixation rates, especially with the stochastic sequence coverage of our dataset (we limited this analysis to sites with ≥ 2X coverage). Still, our set of candidate genes with *d*_N_/*d*_S_-based signatures of positive selection on the *Megaladapis* lineage included the growth hormone receptor (*GHR*), a finding of interest given the large reconstructed body size *M. edwardsi* (~85 kg)^3,8^, one of the ‘giant’ extinct subfossil lemurs. Yet we did not observe an enrichment for body size or growth-related functional pathways among the overall candidate gene set. Given that body size variation is often highly polygenic^73–75^, the absence of such an enrichment is not unexpected.

We have more confidence in the connection of several evolutionary genomic results to potential *Megaladapis* diet-related adaptations. Specifically, we identified enrichments for convergent amino acid evolution between *M. edwardsi* and the golden snub-nosed monkey (a folivore) across genes with hydrolase activity functions, and between *M. edwardsi* and horse (an herbivore) across genes with brush border functions. Hydrolases help to break down plant secondary compounds^43^, while brush border microvilli play crucial roles in nutrient absorption and hydrolysis in the gut ^49,52^. Additionally, our set of candidate genes with *d*_N_/*d*_S_-based signatures of positive selection on the *Megaladapis* lineage included *SULT1C2*, which encodes an enzyme involved in the detoxification of toxic phenolic compounds common in leafy plants^35,37^. In the future, the biology of these *Megaladapis* molecular changes in could be examined via colobine monkey and horse *in vivo* or *in vitro* studies or other functional evolutionary genomics approaches^76^, as appropriate.

Our evolutionary genomic findings support developing reconstructions of *Megaladapis* as a specialized folivore. Specifically, molar microwear patterns are important proxies for inference of the diet of an individual animal in the weeks prior to its death: shearing tough foods, such as leaves, often produces scratches on the tooth surface, while consuming fruits with hard pericarps or hard seeds may lead to punctures or pits^40^. *Megaladapis edwardsi* microwear patterns feature scratches characteristic of leaf-eaters^38,39^, with similarities to extant primate folivores, including *Presbytis entellus*^38^ and *Lepilemur petteri* (with a habitat in Madagascar overlapping that of *M. edwardsi*)^39^. Furthermore, *M. edwardsi* dental topography features including low molar occlusal surface complexity and high “Dirichlet normal energy” (a measurement that roughly captures crown profile “relief” or more precisely, changes in the direction of occlusal surface tangents) suggest adaptation for efficiency in shearing leaves^41^.

Stable isotope data are also consistent with the reconstruction of a folivorous diet for *Megaladapis*. Specifically, carbon isotope ratios can help differentiate the relative contributions of C_3_ plants, C_4_ plants and stem/leaf succulents, as well as non-photosynthetic plant tissues (e.g., fruits, flowers) in the diets of extinct species^60,61^. The spiny thicket habitat in southern and southwest Madagascar is dominated by succulents with patches of C_4_ grasslands and C_3_ trees, yet *M. edwardsi* carbon isotope ratios ubiquitously suggest a C_3_-based, herbivorous leafy diet^60^.

Several *M. edwardsi* craniodental traits suggest adaptations to a “browsing-via-plucking” mode of leaf-eating, including the loss of upper incisors, ventrally flexed nasal bones, posteriorly expanded temporomandibular joint surfaces for compressive mastication, and a post-canine diastema^9,25,38,77^. Similar to koalas, *M. edwardsi* has a caudally positioned foramen magnum and limited mid-face projection relative to length, interpreted to facilitate greater head movement to facilitate direct foraging on leaves^9,77^.

Finally, variation in infraorbital foramen size (IOF) is an osteological proxy for “maxillary mechanoreception” (i.e., how mammals use their “snouts” to acquire and process foods) and dietary inference, at least among primates: Frugivores tend to have larger IOF areas relative to folivores and insectivores, perhaps reflecting adaptations for selecting and evaluating fruit^78^. The relative IOF area of *Megaladapis edwardsi* is significantly smaller than that of any frugivorous extant lemur and is instead more similar to relatively more strict extant lemur folivores (e.g., *Lepilemur*)^59^.

### Conclusion

Overall, our work highlights both the challenges and exciting prospects of non-human primate paleogenomics. Many non-human primates live in the tropics or sub-tropics, which can be challenging environments for ancient DNA recovery and analysis. In this study, we had the opportunity to focus a concerted shotgun sequencing effort on a particular *Megaladapis edwardsi* sample with higher-than-typical levels of ancient DNA preservation for a subfossil specimen from Madagascar, resulting in the first nuclear genome sequence for an extinct non-human primate. In the future, improved methods to extract DNA from tropical and subtropical samples^79^ alongside further technological innovations may facilitate future recoveries of additional nuclear genome sequences from other extinct lemurs or non-lemur primates. For now, we are excited to have been able to analyze *M. edwardsi* nuclear genome sequences for insights into the evolutionary biology and behavioral ecology of this extinct subfossil lemur and to robustly resolve its phylogenetic relationship with other lemurs.

## Data Availability

All sequence data newly generated for this study have been deposited in the Sequence Read Archive for *Megaladapis edwardsi* (PRJNA445550), *Eulemur rufifrons* (PRJNA445550) and *Lepilemur mustelinus* (PRJNA445550). Extant/extinct lemur *mpileup* exon files, masked/unmasked 2X *M. edwardsi* sequences integrated with the UCSC species and extant lemurs alignment data sets, gene alignments used in gene tree phylogeny estimation (input and output files), *d*_N_/*d*_S_ input nucleotide files and resulting output table, functional enrichment output tables and genomic convergence output (supplementary tables) have been deposited to the Dryad Digital Repository (https://doi.org/10.5061/dryad.5qfttdz3c). Code for the *d*_N_/*d*_s_ and genomic convergence analyses used in this manuscript have been made available through the following github repositories: https://github.com/RBankoff/PAML_Scripts/ and https://github.com/MehreenRuhi/conv.

## Methods

### Sample preparation and sequencing of the *Megaladapis edwardsi* nuclear genome

#### DNA extraction

The UA 5180 mandible was sampled under a collaborative agreement with the Department of Paleontology and Biological Anthropology at the University of Antananarivo, Madagascar. All ancient materials were processed in dedicated sterile facilities with positive pressure at the Pennsylvania State University, with physically separate post-PCR processing facilities. As part of a previous study^6^, we identified a *Megaladapis edwardsi* mandible, UA 5180, from the site of Beloha Anavoha, southern Madagascar^6^, with sufficient endogenous DNA quality and quantity for a whole-nuclear genome shotgun sequencing effort. The UA 5180 specimen was directly AMS ^14^C dated (CAMS 142541) as part of a previous study^1^. We have used a new calibration curve (SHCal20)^80^ to recalibrate^81^ the ^14^C age (1640±30) to 1,475 ± 65 cal yr BP. For this study we prepared eight additional DNA extractions from the UA 5180 using an established protocol for animal hard tissue^82^ and following our previously described subsampling strategy^6^.

#### Library preparation

We prepared a total of nine double-stranded libraries with barcoded adapters from specimen UA 5180 suitable for Illumina massively parallel sequencing platforms following the Meyer and Kircher protocol^83^. We used 50μL of template as input for the initial blunt-end repair step without the enzymatic removal of uracil residues and abasic sites. The post-reaction purification steps were carried out using the Qiagen MinElute PCR Purification kit after blunt-end repair and carboxyl-coated magnetic beads (Solid Phase Reversible Immobilization or SPRI) for the adapter ligation and fill-in steps. The final elution volume of 20μL in TET (TE-Tween-20) was then used as template for the indexing reaction. Libraries were barcoded using a single unique P7 index primer where ‘xxxxxxx’ represents the specific barcode for a library (200nM, 5’-CAAGCAGAAGACGGCATACGAGATxxxxxxxGTGACTGGAGTTCAGACGTGT-3’) was added to each library (n=9) with a universal IS4 forward primer (200nM, 5’-AATGATAACGGCGACCGAGATCTACACTCTTTCCCTACACGACGCTCTT-3’)^83^ in a 50μL reaction that also included PCR buffer, 2mM MgSO4, 200 μM dNTPs, and 2.5 U Platinum *Taq* High Fidelity DNA Polymerase (Thermo Scientific) prepared in the ancient DNA facility. Amplification of these libraries was performed under cycling conditions of a 5 min denaturation at 94°C; 24 cycles of 20 sec at 94°C, 15 sec at 60°C, and 20 sec at 68°C; with a final extension of 5 min at 60°C. SPRI beads were used for post-reaction clean-up with elution in 15μL of TET (Tris EDTA-Tween-20) buffer.

#### Sequencing

These nine uniquely indexed libraries were subject to multiple sequence runs at the Pennsylvania State Huck Institutes Genomics Core Facility and at the Schuster lab at Penn State (Illumina HiSeq 2000 and 2500, 75-bp paired-end reads), generating a total of 2,139,275,851 paired-end reads and 328 giga base pairs (Gbp) of sequence data across 15 total lanes. Each lane contained only one library; some libraries were sequenced across multiple lanes. These sequences have been deposited in the NCBI Sequence Read Archive, Accession no. SRP136389 (SRA BioProject no. PRJNA445550).

#### Bioinformatic processing of sequence data

From these raw reads, the forward and reverse adapter sequences (introduced as part of the library preparation protocol) were trimmed, and overlapping paired-end reads were merged using the *MergeReadsFastQ_cc* script^84^ with default settings, using a minimum 11 nucleotide (nt) overlap and a phred quality score of 20 for merged sites. For the unmerged but properly paired reads, Trimmomatic^85^ was used to trim bases downstream of any site with a quality score ≤20, requiring that both unmerged reads pass quality filters to be retained (e.g., minimum read length of 20 bp). Since PCR amplification and sequencing of the same DNA fragment may create identical reads (e.g., the bases in reads are exact duplicates), we used custom perl scripts to collapse such identical reads within each separate library to the single read with the best sum of fastq quality scores before mapping (*SI Appendix*, **Table S1**).

### Sample preparation and sequencing of modern lemur genomes

#### DNA extraction

Ear punches were obtained from wild-caught lemurs, *Lepilemur mustelinus* (Weasel sportive lemur, TVY7.125 from Runhua, Madagascar) and *Eulemur rufifrons* (RANO5.15 from Ranomafana and ISA2.23 from Isalo) with capture and sampling procedures approved by the Institutional Animal Care and Use Committee of Omaha’s Henry Doorly Zoo and Aquarium (#12-101). Collection and export permits were obtained from Madagascar National Parks, and the Ministère de l’Environnement, de l’Ecologie et des Forêts (MEEF) of Madagascar. The samples were imported under requisite CITES permits from the U.S. Fish and Wildlife Service. Genomic DNA was extracted from these blood and tissue samples using a standard phenol/chloroform method from wild-caught individuals (as performed in Kistler et al.^6^) at Penn State University.

#### Library preparation and sequencing

The *L. mustelinus* specimen underwent double-stranded library preparation and indexing following the protocols described above for *M. edwardsi*. The single library generated was shotgun sequenced at the University of California Los Angeles Genomics Center Illumina HiSeq 2500 (100-bp paired-end) (*SI Appendix*, **Table S1**). These sequence reads have been deposited in the NCBI Sequence Read Archive, Accession no. SRP136389 (SRA BioProject no. PRJNA445550). Libraries were prepared for the two *E. rufifrons* specimens with the TruSeq PCR-free library preparation kit, with subsequent whole genome sequencing performed at the HudsonAlpha Institute for Biotechnology (Genomic Services Lab) on the Illumina HiSeq X Ten (150-bp paired-end, one sample per lane) (*SI Appendix*, **Table S1**). The *E. rufifrons* sequence reads have been deposited in the NCBI Sequence Read Archive, Accession no. SRP136389 (SRA BioProject no. PRJNA445550).

#### Existing genomic data

Previously published primate whole genome sequence read data were used for *Daubentonia madagascariensis* (SRA043766.1)^12^, *Propithecus diadema* (PRJNA317769)^14^, *Microcebus murinus* (PRJNA285159)^13^ and *Rhinopithecus roxellana*, (PRJNA230020)^22^.

### Authenticity of *M. edwardsi* genomic data

To assess the authenticity of the *M. edwardsi* ancient nuclear genome sequence data, we considered the fragment length distributions of all sequenced libraries and the nucleotide damage pattern of the mapped reads, and we also estimated the proportion of human DNA contamination.

The fragment length distribution (FLD) for each of the sequenced *M. edwardsi* libraries (n=9; with 5 of the libraries sequenced twice) was composed of abundant short DNA fragments, which is characteristic of ancient specimens^16,86^ (*SI Appendix*, **Figs. S3 and S4**).

The authenticity of our *M. edwardsi* data is supported by the fragment size distribution (*SI Appendix*, **Fig. S5A**), base frequency fragmentation prior to read starts (*SI Appendix*, **Fig. S5B**), and elevated rates of C>T and G>A mismatches as expected at read ends (up to 20%) (*SI Appendix*, **Fig. S5C**). To further characterize DNA damage and degradation, we focused on DNA nucleotide mismatches detectable in double- and single-stranded overhangs (δ¤ and δs, respectively) that due to cytosine deamination are typically over-represented in paleogenome samples in the 5’ termini as cytosine to thymine (C>T) mismatches (guanine to adenine or G>A on the complementary 3’ strand)^18,86^. We used mapDamage 2.0^87^ to quantify post-mortem damage signals in the alignment of *M. edwardsi* reads to the hg19 hard-masked exon reference (UCSC Genome Browser)^88^ from our genomic analyses (outlined below). Through mapDamage analysis, we estimated the probability of cytosine deamination^18^ as *δ*_s_ = 0.73—73% of cytosine residues in single-stranded overhangs have been affected by deamination. To characterize the temporal rate of this chemical damage, we calculated a cytosine deamination rate of 8.55 x 10^−3^ site^−1^ year^−1^, placing deamination in the expected range for bone at a site with an annual mean temperature of 23.42°C^16^. The probability of a nucleotide terminating an overhang was inferred using mapDamage at λ = 0.26 (mean overhang length 3.4nt). Therefore, the first 9nt with the end of a given fragment is expected to contain 95% of misincorporated deoxy-uracil residues under the geometric distribution. Accordingly, for our analyses of the *M. edwardsi* nuclear genome sequence, for all sequence reads we hard-masked (i.e., replaced with ‘N’) sites potentially affected by cytosine deamination (5’ T residues and 3’ A residues)^89^ within 9nt of fragment ends accordingly.

To estimate the level of human DNA contamination in our dataset, we aligned 45 million raw sequence reads sampled from across the multiple ancient DNA libraries to both the *M. edwardsi* mtDNA reference genome sequence (NC_026088.1)^6^ and a human mitochondrial genome sequence (haplotype H6A1; EU256375.1) using bwa *aln*^90^ with seeding disabled (-l 16500) and default mapping parameters (-n 0.01 and -o 2), filtering for a minimum read length of 20nt and minimum mapping quality of 20. We assigned 4,930 non-redundant reads to the *M. edwardsi* reference mtDNA, yielding 22X coverage of the complete mitochondrial genome, versus only 85 reads that mapped to the human reference mtDNA (1.7% of the total mapped reads). Of those 85 human-mapped reads, 25 were in regions strongly conserved across primates including *M. edwardsi*. The remaining 60 reads mapped uniquely the human reference genome. We thus estimate ~1.2% contamination of our *M. edwardsi* sequence data with modern human DNA, consistent with or below reported human contamination rates in other studies^20,21^ and contributing negligibly to our gene sequence reconstructions, especially given our minimum 2X sequence coverage requirements.

We roughly estimated the proportion of endogenous *M. edwardsi* DNA in our sequencing libraries to be 6.13% by computing the number of sequence reads (following merging and removal of identical sequence duplicates that were mapped to hg19 exons and flanks (see below; n=4,824,118) times 27.197 (given that these targets comprise ~3% of the nuclear genome), all divided by the total number of all reads sequenced (n=2,139,275,851).

### Sequence read alignments to human exons

With the sequence read length and coverage restrictions of our ancient DNA data, it was not possible to construct a *de novo* assembly of the *M. edwardsi* nuclear genome. Thus, it was necessary to align our sequence reads to an existing reference genome sequence. We focused on exons, which tend to be relatively conserved across species, thereby aiding mapping and alignment efforts between lemur sequence reads and the human reference genome. We prepared an hg19 reference genome with NCBI Reference Sequences (RefSeq) annotations^91^, with hard-masking so that only the RefSeq exons and 100 nucleotide (nt) flanks to either side of each exonic region were available as alignment targets. The inclusion of the 100 nt flanks helps minimize loss of data from exon ends.

The modern lemur and colobine monkey sequence read data were mapped against this modified hg19 reference using bwa *mem*^90^ with slightly relaxed mismatch penalty (option -B 2). The bwa mem algorithm has higher tolerance for divergent reference sequences given a suitable minimum read length (≥70 bp) (i.e., 2% error for a 100 bp alignment)^90^. The resulting SAM files were converted to BAM using SAMtools^92^ and then used to generate exon consensus sequences using SAMtools *mpileup* (default settings).

For *M. edwardsi*, given the shorter read lengths of this dataset, we instead used a lastZ^93^ alignment procedure. The workflow involved parsing the target sequence into overlapping fragments that were then compared iteratively to the query sequences (individually and sequentially) and filtered by score to remove alignment blocks that did not meet the specified criteria^93^. An extension matrix (Dryad) based on an aye-aye-human whole genome alignment^12^ was used to align curated *Megaladapis edwardi* reads to the prepared hg19 reference, which functioned to modify the scoring scheme to reject or continue along a query sequence to reflect homology with variably diverged sequences. The following command line options were used: *format=general:namel, zstartl, endl, text 1, name2, strand2, zstart2, end2, text2, nucs2, quals2, identity,coverage,continuity −ambiguous=iupac*. The alignment output uses lastZ’s “general format” (e.g., one line per alignment block) and reports the aligned pair of sequences as well as the number of mismatches for that pair^93^. IUPAC (International Union of Pure and Applied Chemistry) ambiguity codes for nucleotides (e.g., N, B, D, H, K, M, R, S, V, W, Y) were treated as completely ambiguous and scored as zero when these substitutions were present^93^.

For reads with more than one viable mapping location (lastZ returns all hits rather than a heuristically optimal hit), we calculated the mean identity (percentage of aligned bases matching the target or query), coverage (percentage of the alignment blocks that cover the entire target or query), and continuity (percentage of the alignment blocks that are not gaps) of each location. Given a 5% minimum difference in this mean between the best and second best hit, we retained the top mapping location. Reads with two or more very similar scores (i.e., less than 5% difference between hg19 exons matching the same region of *M. edwardsi*) on this metric were discarded. The lastZ file was sorted according to genomic coordinates, and then any remaining PCR duplicates with matching start and end position coordinates were discarded by retaining the single read amongst all matches with the greatest sum of fastq quality scores.

We generated a simple positional pileup file from the damage-masked *Megaladapis* read alignment, and we summarized exonic nucleotide positions in the “known canonical” reference gene set from the hg19 assembly (http://hgdownload.soe.ucsc.edu/goldenPath/hg19/multiz46way/). We summarized only sites with a strict consensus among *Megaladapis* reads (leading to higher confidence in authenticity with some loss of heterozygous sites), and we enforced strict positionality to maintain the reading frame of the human reference exons, ignoring indels observed in read data. We likewise generated exon consensus sequences in the modern lemur and colobine monkey read alignments to the hg19 exonic reference, using SAMtools *mpileup* (default settings)^92^ to summarize positional nucleotides. From the pileup files, we summarized exon sequences matching the hg19 “known canonical” gene set enforcing a minimum 2X sequence coverage excluding masked sites, spliced exons to match the full transcripts, and added our *M. edwardsi*, modern lemur, and colobine monkey gene sequences with the remaining sequences, resulting in a 53-way multi species-alignment that we used in our analyses. Because our exon extraction and the 46-way alignment were already forced into the human reading frame, further multiple alignment of gene sequences was not needed.

### Phylogenetic analyses

We used a genome-wide maximum likelihood approach to estimate the phylogenetic placement of *Megaladapis edwardsi*. First, a concatenated gene alignment was constructed comprised of 15 primate species at RefSeq gene loci (described above) where at least 50% of *Megaladapis* sites were represented at minimum 2X coverage after damage masking (n=896 loci; 1.07 Mbp). Using RAxML^23^, we estimated a single unrooted phylogeny from the concatenated alignment without partitioning under the GTR GAMMA model (assuming variable nucleotide frequency changes that are independent for each type of nucleotide)^94^.

Independent phylogenies were also estimated from each gene with at least 20% of sites covered across all lemur sequences (n=12,809) using the same GTR GAMMA model, with 100 bootstrap replicates for each gene tree. Mean bootstrap support across all bipartitions was calculated as a measure of gene tree phylogenetic signal (following Salichos and Rokas, 2013)^24^, with resulting values ranging from 7.69% to 98.85% (median 76.38%; *SI Appendix*, **Fig. S7**). As described previously^24,95^, internode certainty (IC) among gene trees with strong phylogenetic signal provides a robust validation of gene tree support for species tree branching order. We therefore also calculated IC across the concatenated gene tree among bootstrap consensus gene trees with at least 90% mean bootstrap consensus support (n=771 loci) using RAxML^23^.

PhyloPic silhouettes were used in Figure 1A, with the following attributions: *Callithrix jacchus, Papio anubis, Rhinopithecus roxellana, Gorilla gorilla, Tarsius syrichta (Carlito syrichta), Otolemur garnetti (Galagonidae*) all under Public Domain Dedication 1.0 license, https://creativecommons.org/publicdomain/zero/1.0/. *Macaca mulatta* and *Homo sapiens sapiens* under Public Domain Mark 1.0 license https://creativecommons.org/publicdomain/mark/1.0/. Credit to T. Michael Keesey (vectorization) and Tony Hisgett (photography) for *the Pan troglodytes* image, under license https://creativecommons.org/licenses/by/3.0/ (modified opacity). Credit to Gareth Monger for the *Pongo pygmaeus abelii* image, https://creativecommons.org/licenses/by/3.0/ (modified opacity).

### *d*_N_/*d*_S_ analyses

We used the codeml function of PAML^29^ to estimate lineage-specific d✓dS ratios across the phylogeny of the six lemurs in our study plus three non-lemur primates (*Homo sapiens, Gorilla gorilla*, and *Pan troglodytes*). Prior to analysis, all sequences were checked for codon completeness across all nine species and any premature stop codons; violating codons were masked with “N”s (https://github.com/RBankoff/PAML_Scripts/) in accordance with the input requirements of codeml. We restricted our analysis to the set of n=3,342 genes with i) ≥100 intact *Megaladapis* codons in our ≥2X sequence coverage data, ii) ≥100 N sites present and aligned across all nine species in this analysis, and iii) *Megaladapis* lineage *d*_S_ values not more than 2 s.d. greater than the genome-wide average *d*_S_ value.

Based on the PAML codeml results, *d*_N_/*d*_S_ ratios were calculated in two ways: first based on the synonymous substitution rate for an individual gene (*d*_N_/*d*_S_), and second, a *d*_N_/*d*_Sgenome_ ratio based on the genome-wide estimate of *d*_S_^12^. The genome-wide estimate is calculated from the total number of synonymous substitutions across all genes divided by the total number of synonymous sites genome-wide^12^. Inferring a *d*_N_/*d*_Sgenome_ ratio is valuable for branches where the synonymous substitutions may be low or zero for an individual gene^12^.

We used the gene ontology database within g:Profiler (https://biit.cs.ut.ee/gprofiler/gost) to identify any functional category enrichment among the set of genes with *Megaladapis edwardsi* lineage *d*_N_/*d*_Sgenome_ values >1.5, using all 3,342 genes analyzed as background.

### Genome-wide convergent amino acid evolution analyses

We translated our multi-species (n=53) gene sequence alignments (for n=21,520 genes) into amino acid sequences. We then queried each possible individual species (n=35) and clade (n=17) comparison with *M. edwardsi* (using ETE3^96^ to navigate the tree), recording the numbers of analyzable amino acid sites and convergent amino acids for each gene.

A convergent amino acid was defined as having the following properties: i) Both *M. edwardsi* and the distant comparison species/clade to which *M. edwardi* was being compared shared the same amino acid at that position. ii) The sister species to *Megaladapis* (*E. rufifrons*) and a member of the outgroup clade to the *M. edwardsi* and *E. rufifrons* clade (either *M. murinus* or *P. diadema*) shared the same amino acid with each other but not *Megaladapis*. iii) The sister species (or member of the sister clade of species) to the distant comparison species/clade and a member of the outgroup clade to the distant comparison-sister species clade also shared the same amino acid with each other but not with the comparison species/clade.

A site was counted as ‘analyzable’ if the following conditions were met: i) At that site, amino acid information was available for all 6 of the species involved in the particular analysis. ii) E. rufifrons and the outgroup species (e.g. *M. murinus*) had identical amino acids to each other at that position. iii) The sister and outgroup representatives for the distant comparison clade also had identical amino acids to each other at that position.

For the results presented in the main text and figures, we used *M. murinus* as the representative of the outgroup clade to the *M. edwardsi*-*E. rufifrons* grouping because there was a greater number of sites with inferred amino acids for *M. murinus* than for *P. diadema*. We repeated all analyses using *P. diadema* to confirm consistency.

For each comparison, we summed the numbers of convergent and analyzable sites across all genes represented in each GO term^42^. For GO categories with ≥ 5 convergent amino acids we tested whether the proportion of convergent sites was significantly different than expected based on the genome-wide ratio. Specifically, for each qualifying category we used a Fisher’s exact test to compare the ratio of convergent to analyzable amino acid positions within that GO category to this ratio for all genes in the genome apart from those included in that category. We computed false discovery rate (FDR)^97^ from the resulting p-values to account for the multiple tests. The code for this analysis is available at https://github.com/MehreenRuhi/conv and the results have been deposited to Dryad (https://doi.org/10.5061/dryad.5qfttdz3c).

## Acknowledgments

We thank the Laboratoire de Primatologie et de Paléontologie des Vertébrés and the Mention Anthropobiologie et Développement Durable (ADD) at the University of Antananarivo for permission to sample the UA 5180 *M. edwardsi* specimen, and Madagascar National Parks and the Ministère de l’Environnement, de l’Ecologie et des Forêts (MEEF) for permission to collect the modern lemur samples included in the study. Different components of this work were supported the Penn State College of the Liberal Arts, the Penn State Huck Institutes of the Life Sciences, and by grants from the National Science Foundation (BCS-1317163 to G.H.P; BCS-1554834 to G.H.P.; BCS-1750598 to L.R.G.; BCS-1749676 to B.E.C.) and from the Ahmanson Foundation (to E.E.L.). We thank Webb Miller for bioinformatic discussions and contributions. Joel Borgerson created the watercolor illustrations of extant and extinct lemurs shown in Figure 1. Computations for this research were performed on the Pennsylvania State University’s Institute for Computational and Data Sciences’ supercomputing cluster. This content is solely the responsibility of the authors and does not necessarily represent the views of the Institute for Computational and Data Sciences.

## Supplementary Materials

**Supplementary Figure 1:** Sequencing depth for *Megaladapis edwardsi* and extant lemur libraries

**Supplementary Figure 2:** Pre- and post-processed read counts for *Megaladapis edwardsi* libraries

**Supplementary Figure 3:** Individual read length distributions for *Megaladapis edwardsi* libraries

**Supplementary Figure 4:** *Megaladapis edwardsi* unique read length distributions

**Supplementary Figure 5:** Read length distribution, DNA fragmentation patterns and frequency of nucleotide misincorporation for a subset of *Megaladapis edwardsi* sequence reads

**Supplementary Figure 6:** Proportion of sites reconstructed per gene in the 2X coverage *Megaladapis edwardsi* damage-masked and damage-unmasked nucleotide dataset

**Supplementary Figure 7:** Mean bootstrap support across independent gene trees

**Supplementary Figure 8:** Lineage-specific *d*_N_/*d*_S_ ratios for *GHR* and *SULT1C2* (full alignment)

**Supplementary Figure 9:** Phylogenetic relationships of species used in genomic convergence analyses

**Supplementary Figure 10:** *Megaladapis edwardsi* genomic convergence results with *R. roxellana* and *E. equus caballus* (*P. diadema* outgroup)

**Supplementary Table 1:** Sequence metrics and accession information

**Supplementary Table 2:** Proportion of genes covered in the 2X damage-masked and nomasked *Megaladapis edwardsi* data set

