## Supplementary Materials for "Evolutionary and phylogenetic insights from a nuclear genome sequence of the extinct, giant ‘subfossil’ koala lemur *Megaladapis edwardsi*"

**Supplementary Figure 1:** Sequencing depth for *Megaladapis edwardsi* and extant lemur libraries

**Supplementary Figure 2:** Pre- and post-processed read counts for *Megaladapis edwardsi* libraries

**Supplementary Figure 3:** Individual read length distributions for *Megaladapis edwardsi* libraries

**Supplementary Figure 4:** *Megaladapis edwardsi* unique read length distributions

**Supplementary Figure 5:** Read length distribution, DNA fragmentation patterns and frequency of nucleotide misincorporation for a subset of *Megaladapis edwardsi* sequence reads

**Supplementary Figure 6:** Proportion of sites reconstructed per gene in the 2X coverage *Megaladapis edwardsi* damage-masked and damage-unmasked nucleotide dataset

**Supplementary Figure 7:** Mean bootstrap support across independent gene trees

**Supplementary Figure 8:** Lineage-specific $d_N/d_S$ ratios for *GHR* and *SULT1C2* (full alignment)

**Supplementary Figure 9:** Phylogenetic relationships of species used in genomic convergence analyses

**Supplementary Figure 10:** *Megaladapis edwardsi* genomic convergence results with *R. roxellana* and *E. equus caballus* (*P. diadema* outgroup)

**Supplementary Table 1:** Sequence metrics and accession information

**Supplementary Table 2:** Proportion of genes covered in the 2X damage-masked and no-masked *Megaladapis edwardsi* data set

#### Figure S1. Sequencing depth for *Megaladapis edwardsi* and extant lemur libraries

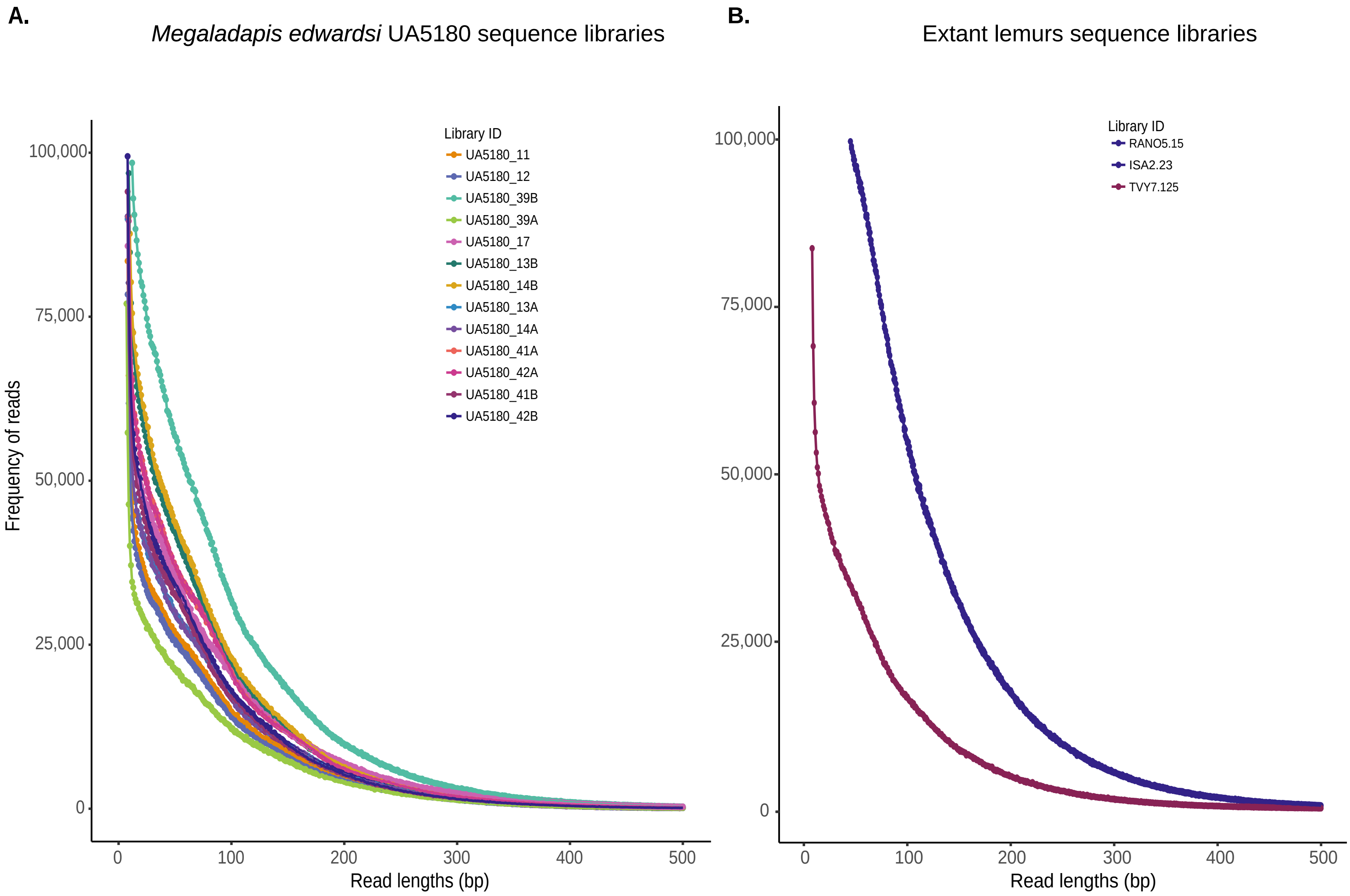


**Figure S1**. Sequencing depth of the total reads generated for A) *Megaladapis edwardsi* libraries (n=9, A and B indicate libraries sequenced twice), and B) extant lemurs, *Eulemur rufifrons* (RANO5.15 and ISA2.23) and *Lepilemur mustelinus* (TVY7.125).

**Figure S2. Pre- and post-processed read counts for *Megaladapis edwardsi* libraries**

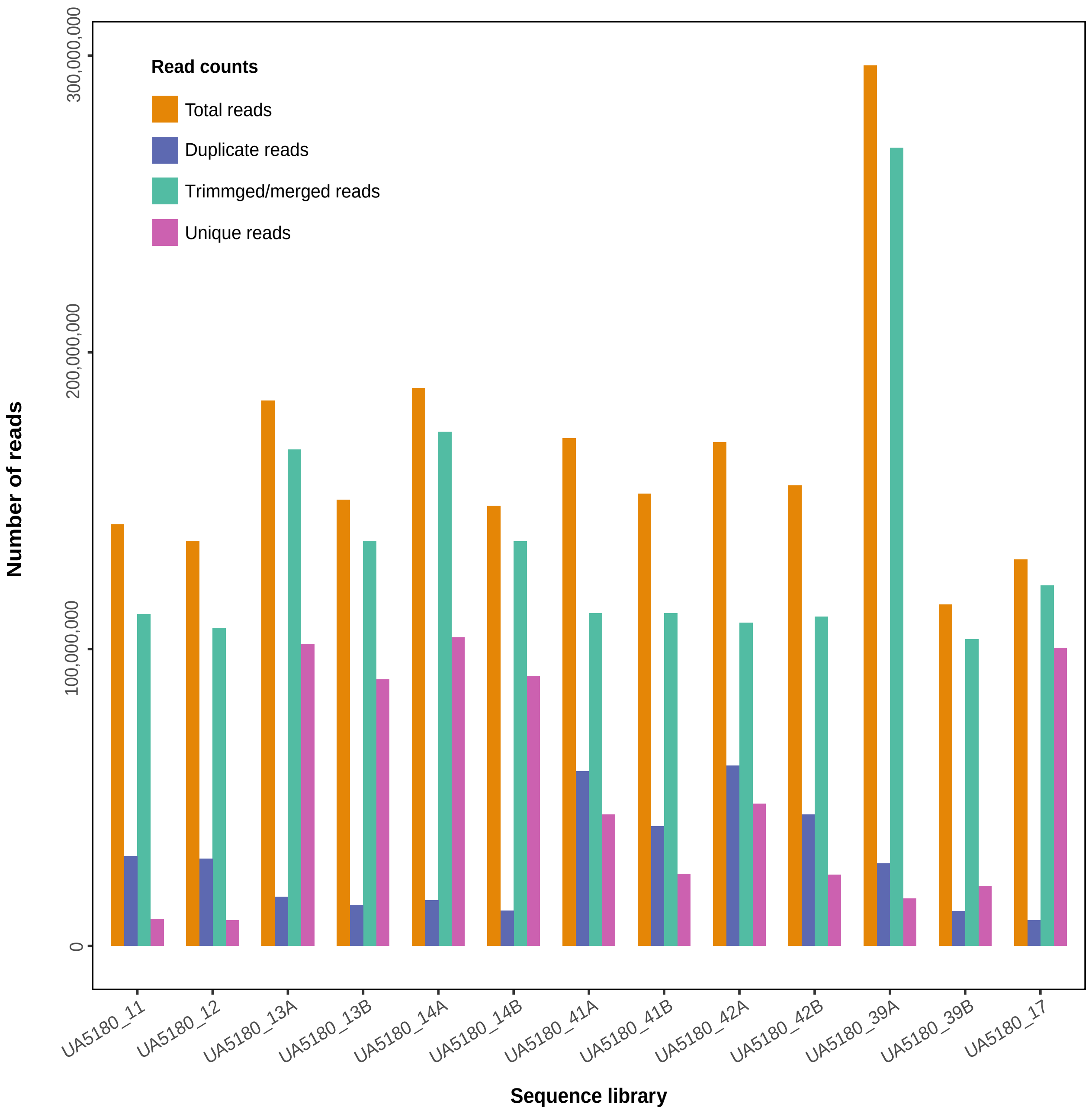


**Figure S2**. The total counts of pre- and post-processed reads for each *Megaladapis edwardsi* library prepared from sample UA5180. “Total reads” are the raw unprocessed sequence reads; “Duplicate reads” are redundant reads based on the number of trimmed and merged reads per sequencing library; “Trimmed/merged reads” are adaptor-trimmed and merged paired-end reads; “Unique reads” are reads filtered to a minimum of 20 bp and base quality of 20.

#### Figure S3. Individual read length distributions for *Megaladapis edwardsi* libraries

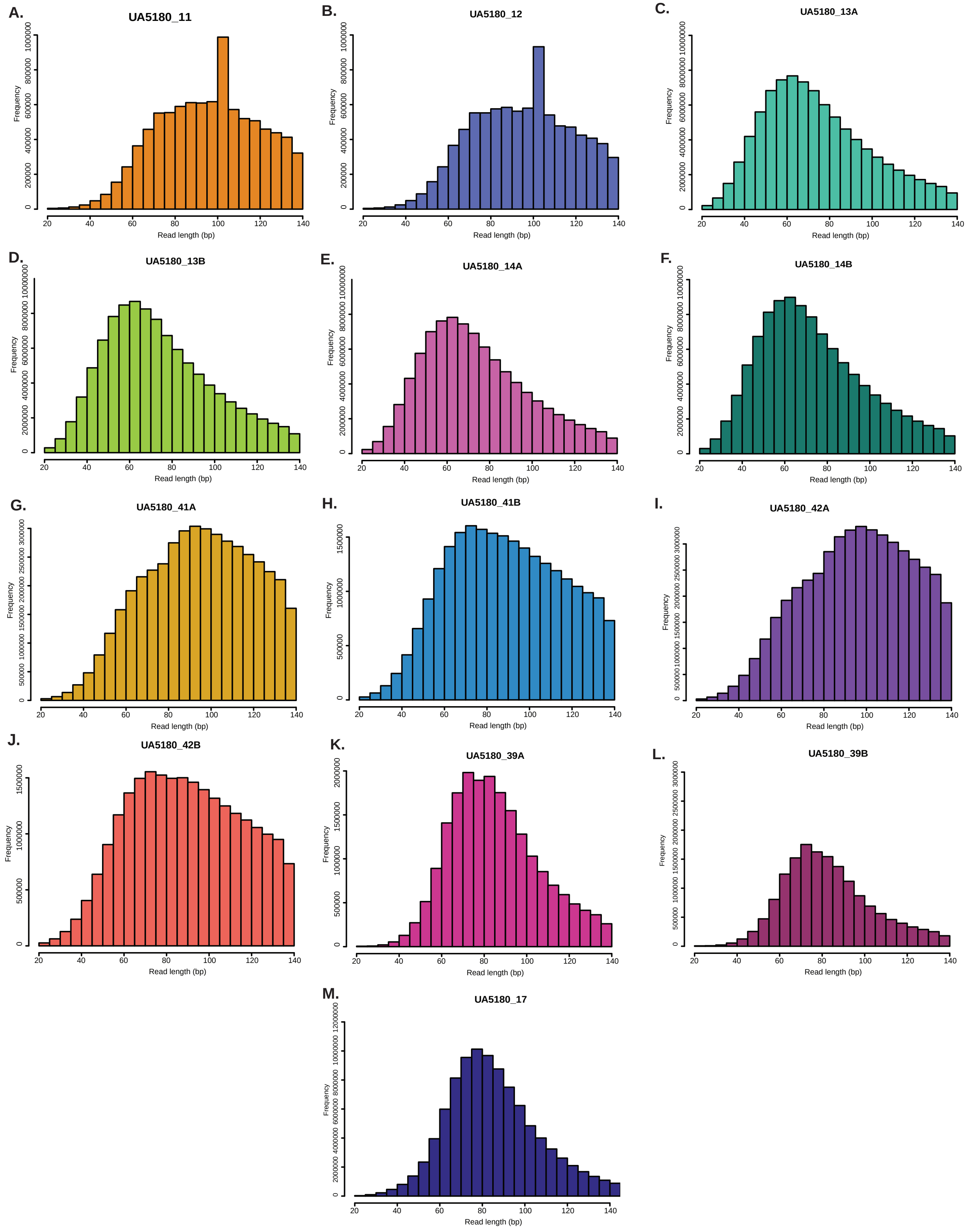


**Figure S3**. Read length distributions (A to M) for quality filtered reads (min. 20bp and quality 20) in each separately sequenced UA5180 *M. edwardsi* library (n=9 libraries, A and B indicate those libraries sequenced twice).

**Figure S4. *Megaladapis edwardsi* unique read length distributions**

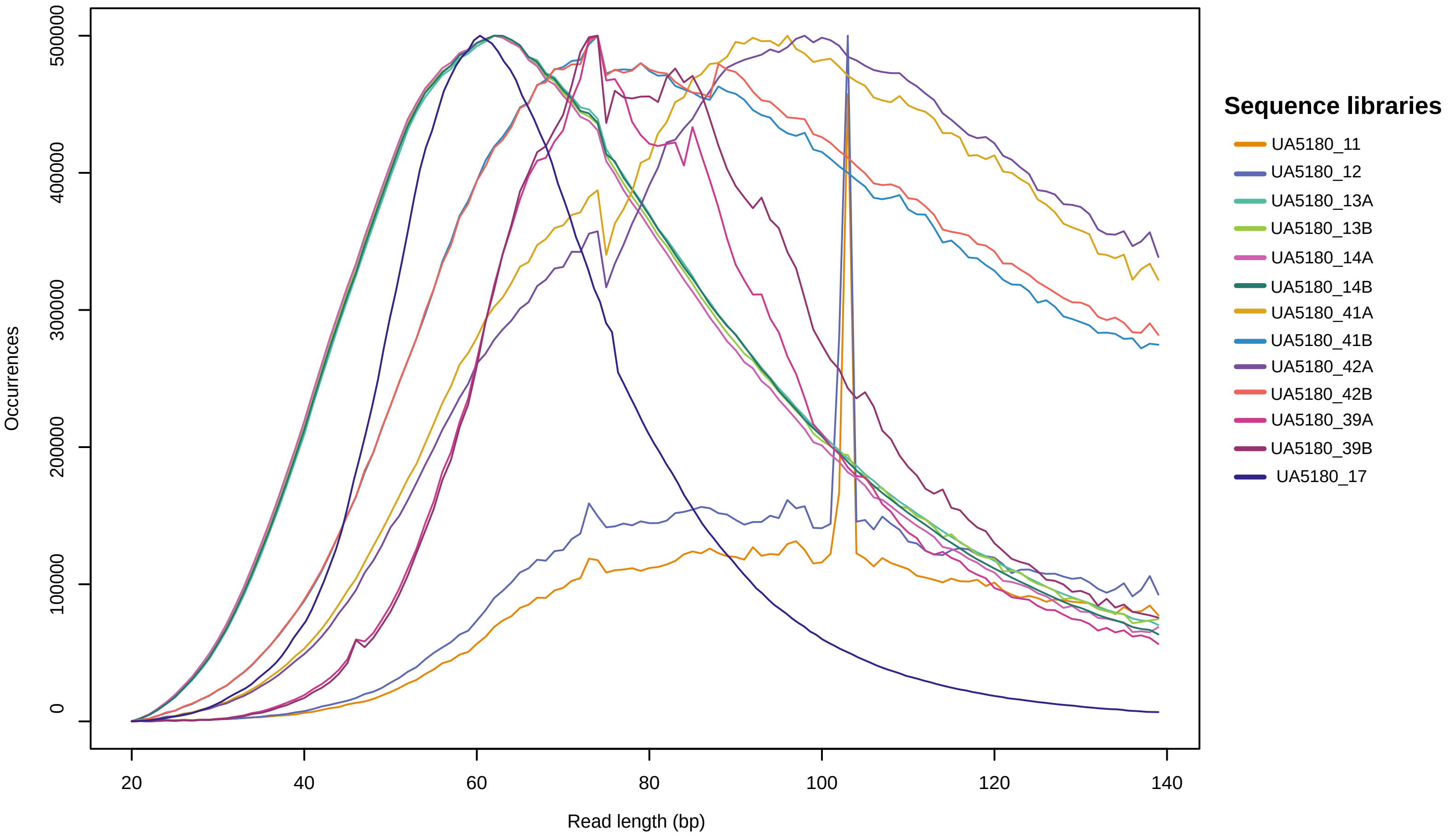


**Figure S4**. Read length overlay of the number of unique reads with a minimum 20bp and quality 20 retained after adapter trimming and merging for each of the 13 sequenced *M. edwardsi* libraries.

**Figure S5. Read length distribution, DNA fragmentation patterns and frequency of nucleotide misincorporation for a subset of *Megaladapis edwardsi* sequence reads**

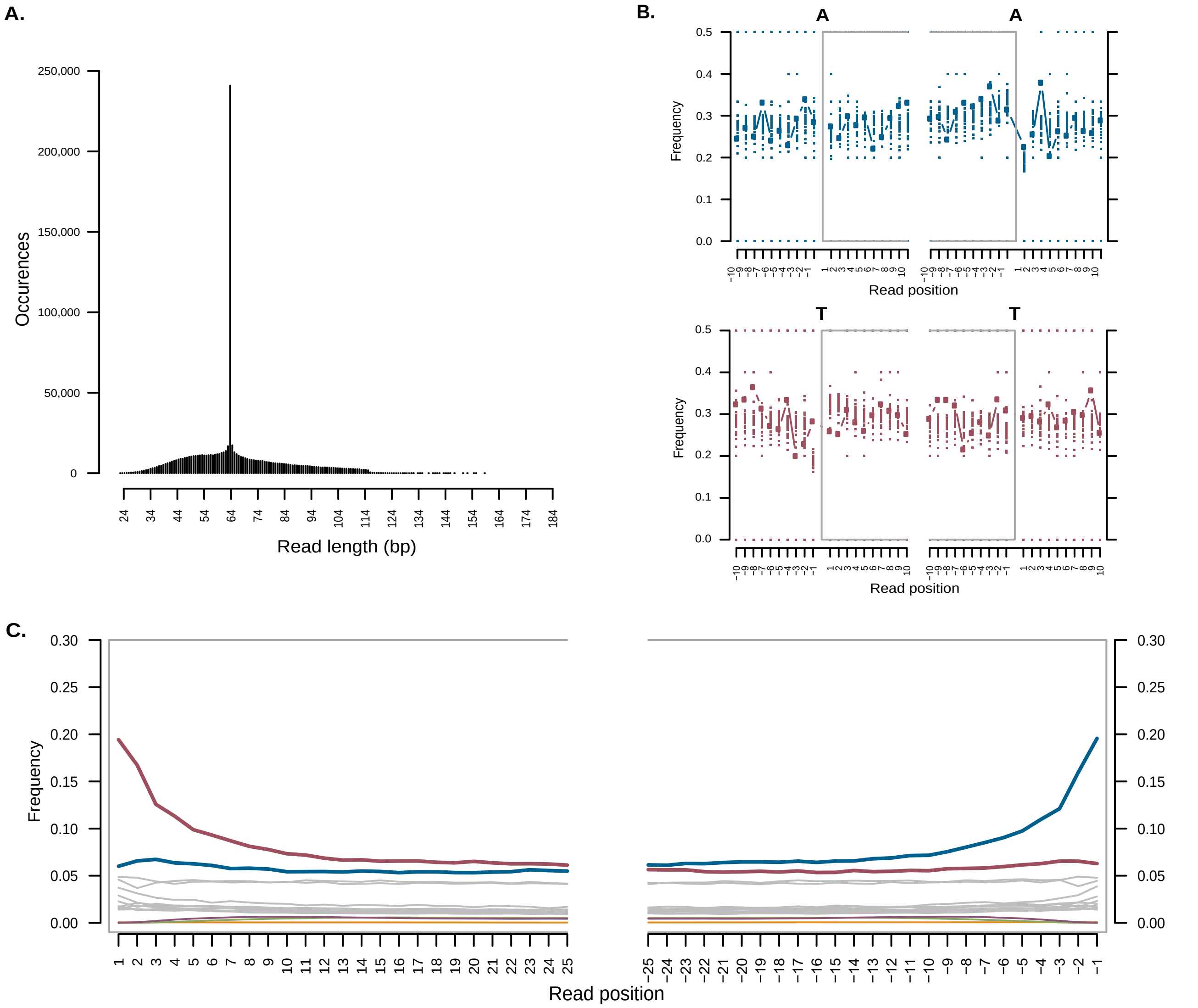


**Figure S5**. Estimates of post-mortem damage for *Megaladapis edwardsi* were generated from a BAM file of the lastZ alignment mapped to a modified version of hg19 (non-exons masked) using mapDamage. A) Histogram of single-end read length distribution. Note, the peak at 64 bp in the fragment size distribution is due to the presence of untrimmed 64 bp single-stranded reads remaining after trimming and merging with an 11 bp overlap. B) Base frequency DNA fragmentation patterns within the read (contained in the grey box) and outside of the read. C) The frequency of nucleotide misincorporation patterns are indicated from the 5' start (left, positive read position numbers) and 3' end (right, negative read position numbers), showing position-specific substitutions: G>A (maroon), C>T (teal), and all other substitutions (grey).

#### Figure S6. Proportion of sites reconstructed per gene in the 2X coverage *Megaladapis edwardsi* damage-masked and damage-unmasked nucleotide dataset

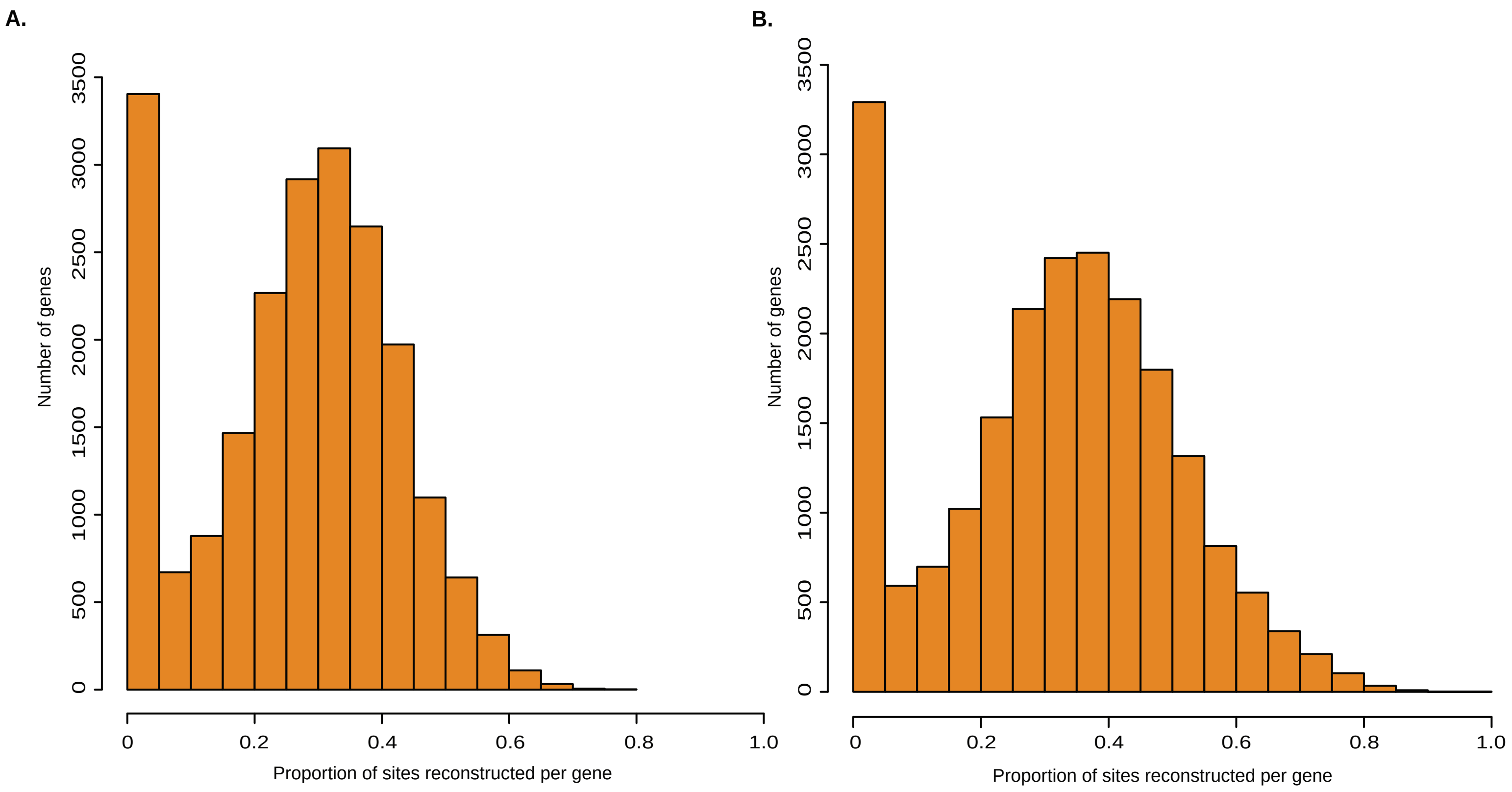


**Figure S6**. *Megaladapis edwardsi* genes were called in damage-masked and damage-unmasked 2X coverage exon sets, containing 21,519 genes total. Genes were called at: A) a median of 28.60% of sites in the damage-masked (17,448 genes called at greater than 10% of sites) and, B) median of 33.11% in the damage-unmasked (17,636 genes called at greater than 10% of sites).

#### Figure S7. Mean bootstrap support across independent gene trees

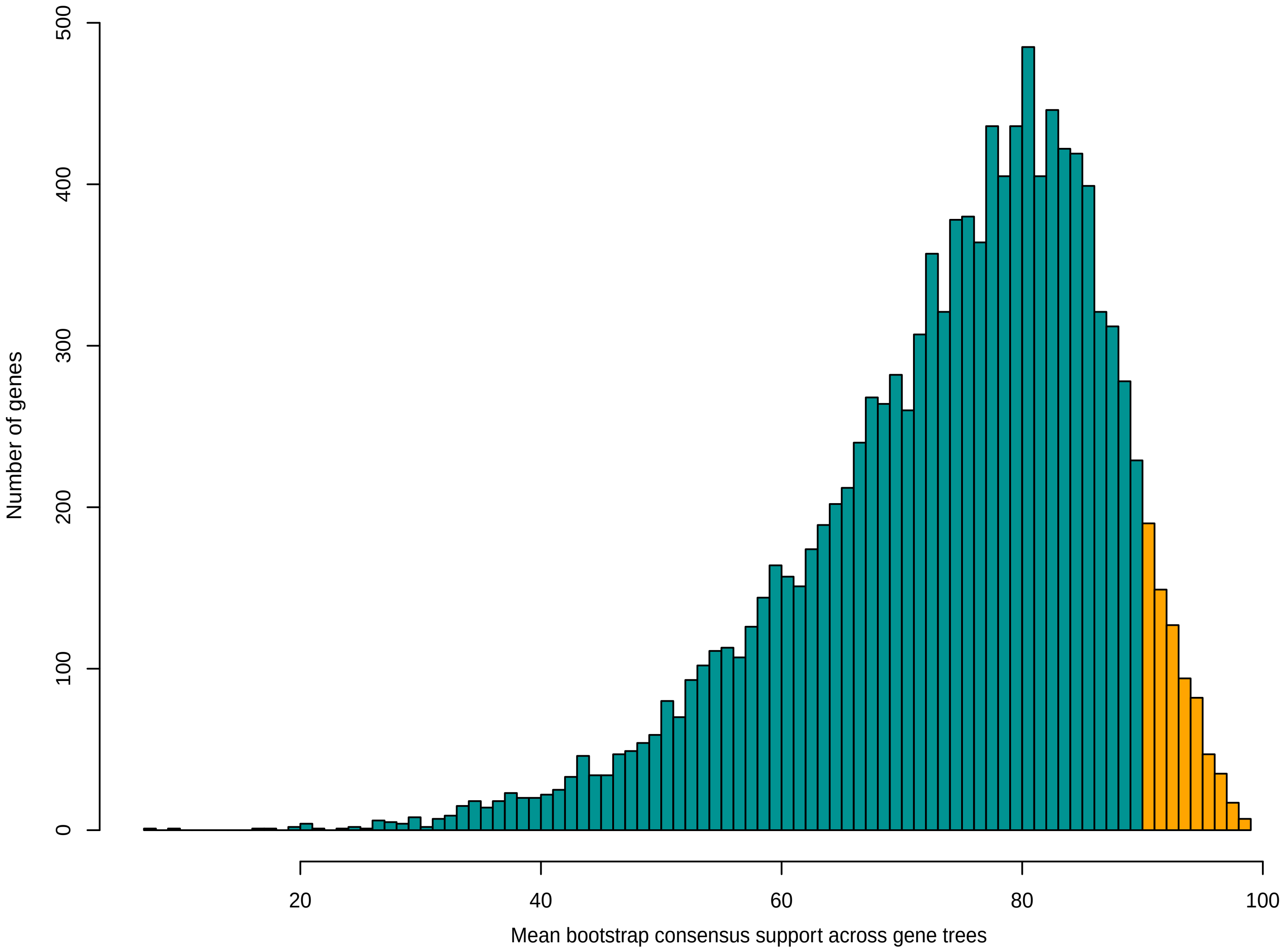


**Figure S7**. Using a database of 12,809 genes with aligned nucleotides present across at least 20% of sites, we estimated independent gene trees (100 bootstrap replicates) using the same model as the full species tree in Figure 1A. The mean level of bootstrap support across all branch bipartitions was calculated as a measure of gene tree phylogenetic signal. The overall average mean bootstrap support value was 74.10% (s.d. = 12.65%; range = 7.69% to 98.85%). Highlighted in orange is the mean bootstrap support that exceeds 90%, representing 771 “strong phylogenetic signal” gene trees.

### Figure S8. Lineage-specific $d_N/d_S$ ratios for *GHR* and *SULT1C2* (full alignment)

**A.**

*GHR* lineage-specific $d_N/d_{Sgenome}$ ratios

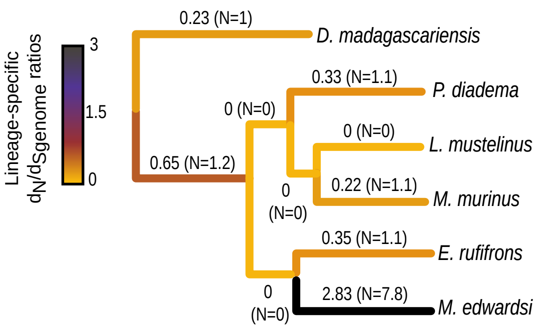


GHR variable amino acid positions

| | 5 | 6 | 8 | 9 | 10 | 41 | 43 | 45 | 62 | 63 | 70 | 71 | 72 | 73 | 74 | 76 | 77 | 78 | 80 | 81 |
|---|---|---|---|---|---|---|---|---|---|---|---|---|---|---|---|---|---|---|---|---|
| *D. madagascariensis* | Q | L | L | T | L | P | L | T | E | T | D | G | V | H | H | L | K | S | G | S |
| *P. diadema* | . | . | . | . | . | . | . | . | . | . | . | . | . | . | . | . | . | **H** | . | . |
| *L. mustelinus* | . | . | . | . | . | . | . | . | . | . | . | . | . | . | . | . | . | **H** | . | . |
| *M. murinus* | . | . | . | . | . | . | . | . | . | . | . | **E** | . | . | . | . | . | **H** | . | . |
| *E. rufifrons* | . | . | . | . | . | . | . | . | . | . | . | . | . | . | . | . | . | **H** | . | . |
| *M. edwardsi* | . | . | . | . | . | . | . | . | . | . | . | **E** | . | . | . | **T** | . | **N** | . | **P** |

| | 82 | 83 | 84 | 85 | 86 | 87 | 153 | 154 | 155 | 157 | 162 | 163 | 164 | 166 | 168 | 169 | 170 | 171 | 172 | 173 |
|---|---|---|---|---|---|---|---|---|---|---|---|---|---|---|---|---|---|---|---|---|
| *D. madagascariensis* | V | Q | L | F | Y | I | I | G | L | W | I | S | L | G | H | A | D | I | Q | V |
| *P. diadema* | . | . | . | . | . | . | . | . | . | . | . | . | . | . | . | . | . | . | . | . |
| *L. mustelinus* | . | . | . | . | . | . | . | . | . | . | . | . | . | . | . | . | . | . | . | . |
| *M. murinus* | . | . | . | . | . | . | . | . | . | . | . | . | . | . | . | . | . | . | . | . |
| *E. rufifrons* | . | . | . | . | . | . | . | **S** | . | . | . | . | . | . | . | . | . | . | . | . |
| *M. edwardsi* | **I** | . | . | . | . | **T** | . | . | . | . | . | . | . | . | . | . | . | . | . | . |

| | 186 | 187 | 207 | 210 | 211 | 234 | 235 | 236 | 237 | 238 | 239 | 240 | 243 | 249 | 251 | 252 |
|---|---|---|---|---|---|---|---|---|---|---|---|---|---|---|---|---|
| *D. madagascariensis* | G | W | M | V | L | Q | R | N | S | E | K | Y | F | V | L | P |
| *P. diadema* | . | . | . | . | . | . | . | . | . | . | . | . | . | **I** | . | . |
| *L. mustelinus* | . | . | . | . | . | . | . | . | . | . | . | . | . | . | . | . |
| *M. murinus* | . | . | . | . | . | . | . | . | . | . | . | . | . | . | . | . |
| *E. rufifrons* | . | . | . | . | . | . | . | . | . | . | . | . | . | . | . | . |
| *M. edwardsi* | . | . | . | . | . | . | . | . | . | . | . | . | . | . | . | . |

**B.**

*SULT1C2* lineage-specific $d_N/d_{Sgenome}$ ratios

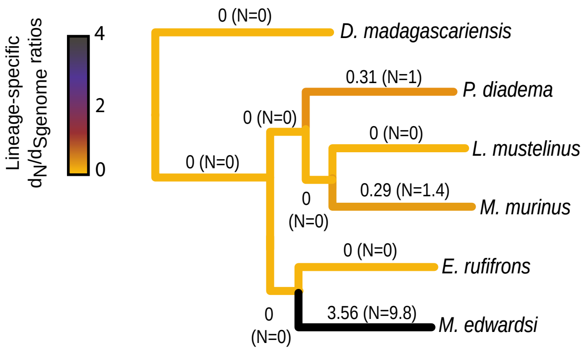


SULT1C2 variable amino acid positions

| | 38 | 39 | 40 | 41 | 42 | 43 | 45 | 46 | 47 | 54 | 56 | 59 | 61 | 66 | 67 | 68 | 70 | 72 | 74 | 138 |
|---|---|---|---|---|---|---|---|---|---|---|---|---|---|---|---|---|---|---|---|---|
| *D. madagascariensis* | K | P | D | D | L | L | C | T | Y | W | Q | V | M | G | D | V | K | Q | A | L |
| *P. diadema* | . | . | . | . | . | . | . | . | . | . | . | . | . | . | . | **M** | . | . | . | . |
| *L. mustelinus* | . | . | . | . | . | . | . | . | . | . | . | . | . | . | . | . | . | . | . | . |
| *M. murinus* | . | . | . | . | . | . | . | . | . | . | . | . | . | . | . | . | . | . | . | . |
| *E. rufifrons* | . | . | . | . | . | . | . | . | . | . | . | . | . | . | . | . | . | . | . | . |
| *M. edwardsi* | . | . | **E** | **H** | **V** | . | **Y** | . | . | . | . | . | . | . | . | **L** | . | . | . | . |

| | 143 | 145 | 154 | 155 | 162 | 164 | 177 | 186 | 235 | 237 | 238 | 239 | 243 | 244 | 248 | 249 | 250 | 254 | 255 | 259 |
|---|---|---|---|---|---|---|---|---|---|---|---|---|---|---|---|---|---|---|---|---|
| *D. madagascariensis* | N | K | D | Q | P | P | G | D | I | Q | E | T | K | M | P | M | T | T | I | I |
| *P. diadema* | . | . | . | . | . | . | . | . | . | . | . | . | . | . | . | . | . | . | . | . |
| *L. mustelinus* | . | . | . | . | . | . | . | . | . | . | . | . | . | . | . | . | . | . | . | . |
| *M. murinus* | . | . | . | . | . | . | . | . | . | . | . | . | . | . | . | . | . | . | . | . |
| *E. rufifrons* | . | . | . | . | . | . | . | . | . | . | . | . | . | . | . | . | . | . | . | . |
| *M. edwardsi* | . | . | . | . | . | . | . | . | . | . | . | . | . | **I** | . | . | **M** | **A** | . | . |

| | 260 | 263 | 265 | 283 | 286 | 287 | 289 | 291 |
|---|---|---|---|---|---|---|---|---|
| *D. madagascariensis* | L | S | S | A | E | R | E | I |
| *P. diadema* | . | . | . | . | . | . | . | . |
| *L. mustelinus* | . | . | . | . | . | . | . | . |
| *M. murinus* | . | . | . | . | . | I | . | . |
| *E. rufifrons* | . | . | . | . | . | . | . | . |
| *M. edwardsi* | **V** | . | . | . | . | . | . | . |

**Figure S8.** Using a maximum likelihood approach implemented in PAML, Lineage-specific ratios of the rates (*d*) of nonsynonymous (N) vs. synonymous (S) substitution along ancestral and terminal branches estimated with a maximum likelihood-based approach for A) the growth hormone receptor (*GHR*) and B) sulfotransferase 1C2 (*SULT1C2*) genes. For each branch, the $d_S$ denominator is based on the genome-wide synonymous substitution rate. $d_N/d_{Sgenome}$ estimates are recorded next to each branch and depicted by the heatmap. The estimated number of N substitutions for each branch are reported within the parentheses. Branch lengths shown are based on those from Figure 1A rather than these individual genes. For each gene, alignments of inferred amino acid residues for the encoded proteins are shown for all variable positions. Amino acid residues identical to those for *D. madagascariensis* are depicted with "." and amino acid position numbers are based on the human reference sequence (hg19/GRCh37).

#### Figure S9. Phylogenetic relationships of species used in genomic convergence analyses

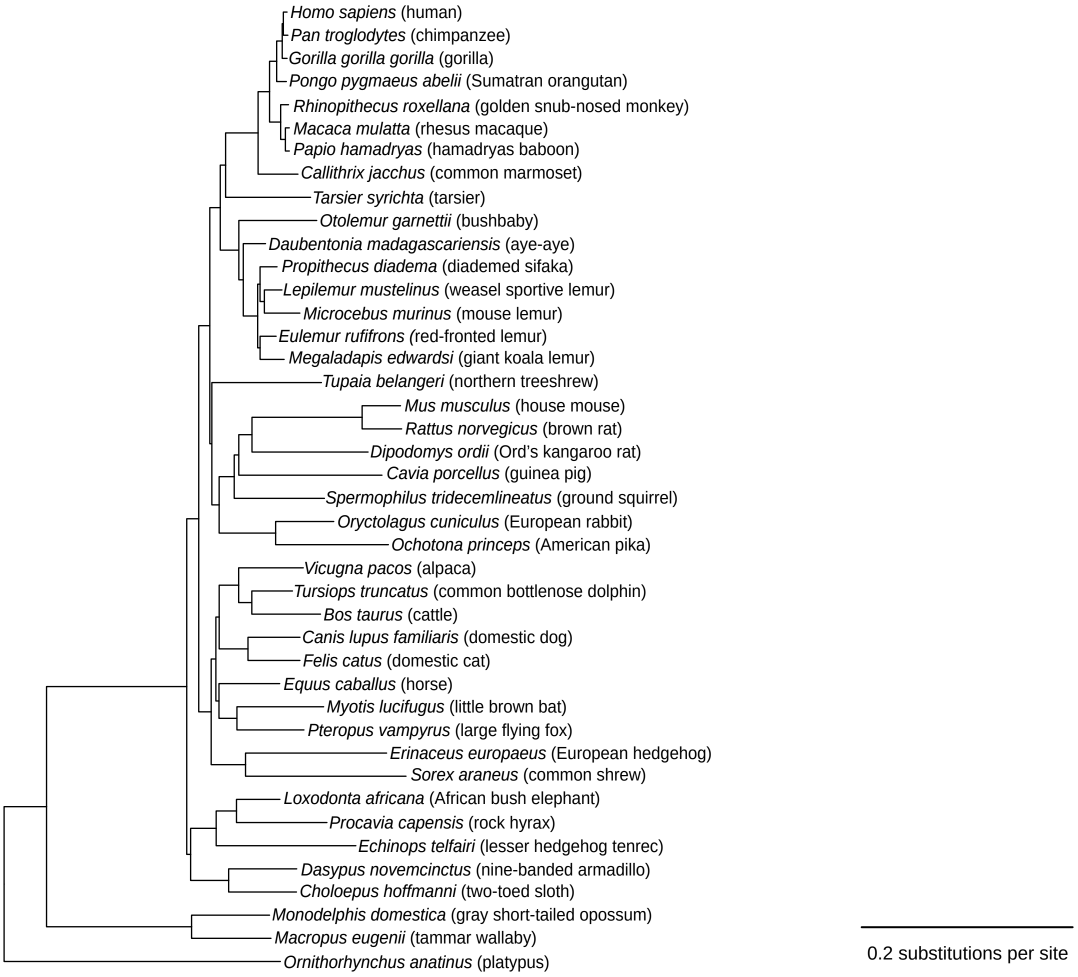


**Figure S9**. Phylogenetic representation of the species from the 46-way UCSC vertebrate alignment joined with the newly sequenced *M. edwardsi*, extant lemurs and colobine. For the convergence analyses, the nucleotide alignments were translated into amino acids, and queried for all possible convergences between species.

#### Figure S10. *Megaladapis edwardsi* genomic convergence results with *R. roxellana* and *E. equus caballus* (*P. diadema* outgroup)

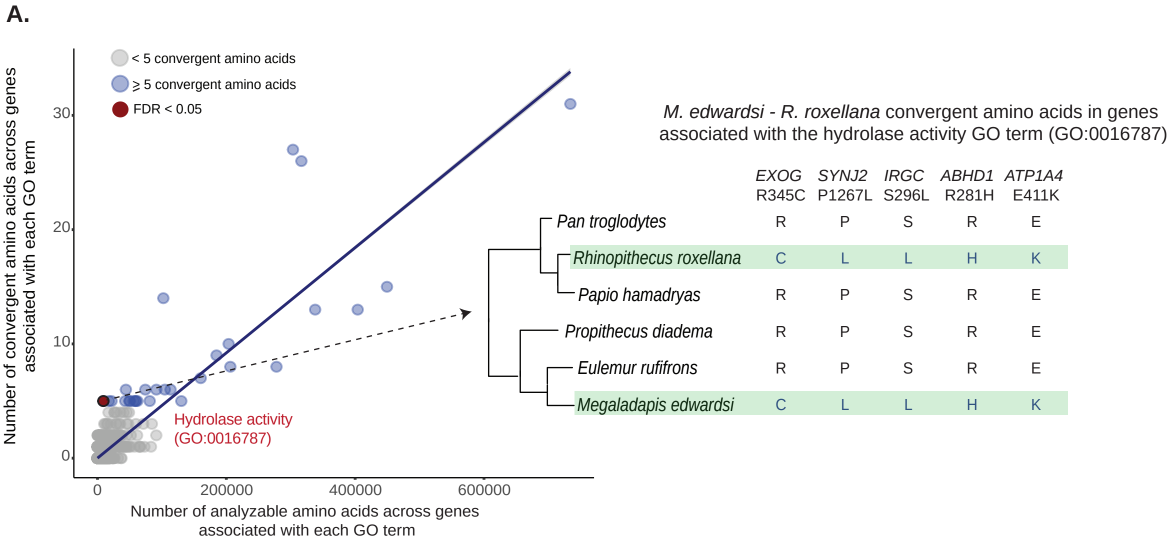


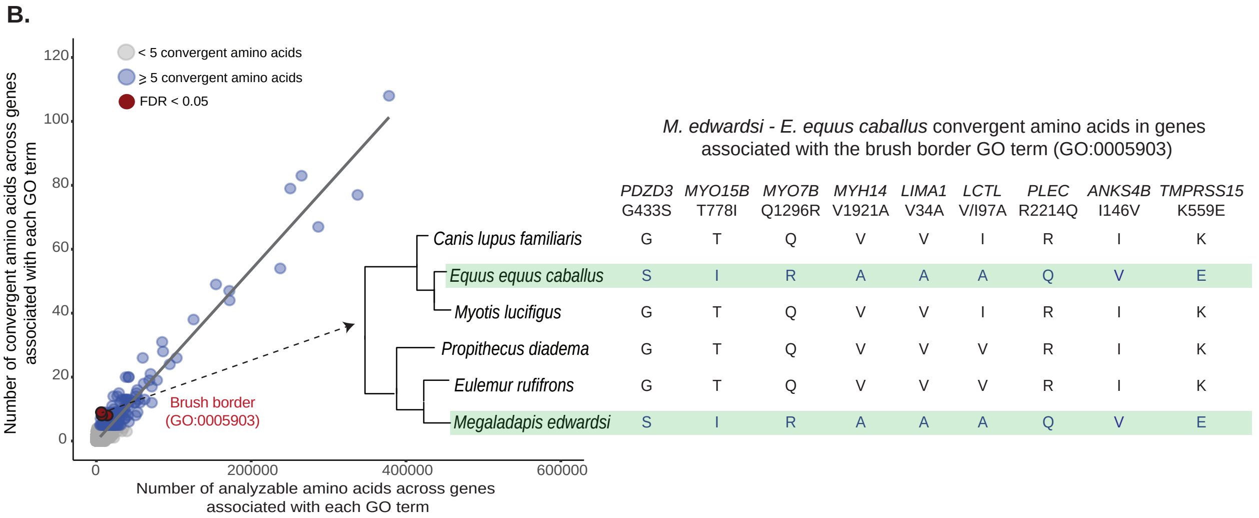


**Figure S10**. Results from scans to identify Gene Ontology (GO) functional categories with unusual proportions (relative to genome-wide expectations) of inferred convergent amino acid positions between A) *M. edwardsi* and the folivore *R. roxellana* and B) *M. edwardsi* and the herbivore *E. equus caballus*. Convergent positions are those with identical residues between *M. edwardsi* and the comparison species, but for which the sister and an outgroup species (for each of the comparison species) share a distinct amino acid residue (shown at right). At left, the number of analyzable amino acid positions and convergent amino acid positions for each GO term. For terms with ≥ 5 convergent amino acids we tested whether the proportion of convergent sites was significantly different than expected based on the genome-wide ratio and computed false discovery rates (FDR) to account for the multiple tests. For two highlighted GO terms, all convergent amino acid positions between *M. edwardsi* and the comparison species along with gene name and position (based on the human reference sequence) are shown.

#### Table S1. Sequence metrics and accession information *Megaladapis edwardsi* shotgun sequenced libraries

| Taxon | Sample ID | Site | Tissue | Cal Yrs. BP | Library ID | Master ID | BioProject Accession | Total reads | Merged reads* | Duplicate reads** (%) | Unique reads*** | % Unique reads |
|---|---|---|---|---|---|---|---|---|---|---|---|---|
| | | | | | Bar13_Dec | UA5180_13A | SRX3843220 | 183,815,699 | 167,293,010 | 8.99 | 101,759,510 | 55.36 |
| | | | | | Bar13_June | UA5180_13B | SRX3843208 | 150,289,729 | 136,466,616 | 9.2 | 89,752,467 | 59.72 |
| | | | | | Bar14_Dec | UA5180_14A | SRX3843223 | 187,934,336 | 173,244,552 | 8.18 | 103,995,978 | 55.34 |
| | | | | | Bar14_June | UA5180_14B | SRX3843209 | 148,344,892 | 136,385,993 | 8.06 | 90,997,592 | 61.34 |
| | | | | | Bar41_July | UA5180_41A | SRX3843212 | 171,073,138 | 112,145,675 | 34.45 | 44,267,729 | 25.88 |
| | | | | | Bar41_Nov | UA5180_41B | SRX3843211 | 152,387,056 | 112,120,011 | 26.42 | 24,303,403 | 15.95 |
| *Megaladapis edwardsi* | UA5180 | Anavoha | Mandible | 1546-1410 | Bar42_July | UA5180_42A | SRX3843213 | 169,686,911 | 108,941,131 | 35.8 | 47,878,427 | 28.22 |
| | | | | | Bar42_Nov | UA5180_42B | SRX3843210 | 155,204,251 | 110,990,589 | 28.49 | 23,972,353 | 15.45 |
| | | | | | Bar11 | UA5180_11 | SRX3843221 | 142,059,057 | 111,797,291 | 21.3 | 9,151,117 | 6.44 |
| | | | | | Bar12 | UA5180_12 | SRX3843222 | 136,491,202 | 107,080,700 | 21.55 | 8,743,143 | 6.41 |
| | | | | | PJX039_June | UA5180_39A | SRX3843218 | 296,737,001 | 268,882,310 | 9.39 | 15,958,787 | 5.38 |
| | | | | | PJX039_May | UA5180_39B | SRX3843219 | 115,052,998 | 103,300,322 | 10.22 | 20,144,369 | 17.51 |
| | | | | | Schuster | UA5180_17 | SRX3843216 | 130,199,581 | 121,515,872 | 6.67 | 100,489,937 | 77.18 |

*Merged reads are adaptor-trimmed and merged paired-end reads, without duplicates removed
**Duplicate reads are redundant reads based on the number of trimmed and merged reads relative to the total reads sequenced
***Unique reads are reads filtered to min 20bp, quality 20 and duplicates removed

**Extant lemurs shotgun sequenced libraries**

| Taxon | Sample ID | Site | Tissue | Date | Library ID | BioProject Accession | Total reads | Unique reads* | Average depth** |
|---|---|---|---|---|---|---|---|---|---|
| *Eulemur rufifrons* | RANO5.15 | Ranomafana | Ear punch | ~2016 | Eulemur_S1 | SRX3843217 | 393,469,627 | 5,279,482 | 7.55 (+/- 7.02) |
| | ISA2.23 | Isalo | Ear punch | ~2016 | Eulemur_S2 | SRX3843214 | 392,627,766 | | |
| *Lepilemur mustelinus* | TVY7.125 | Runhua | Ear punch | ~2016 | Lepilemur | SRX3843215 | 172,058,967 | 6,145,696 | 6.40 (+/-5.85) |

*Unique reads are reads remaining after adapter trimming and read merging, filtered to min 20bp
**Average depth calculated using SAMtools depth

**Accessions for lemurs and colobine used in analyses**

| Taxon | BioProject Accession |
|---|---|
| *Daubentonia madagascariensis* | SRA043766.1 |
| *Propithecus diadema* | PRJNA317769 |
| *Rhinopithecus roxellana* | PRJNA230020 |
| *Microcebus murinus* | PRJNA285159 |

#### Table S2. Proportion of genes covered in the 2X damage-masked and no-masked *Megaladapis edwardsi* data set

**2X damage-masked**

| % of sites at 2X | # of genes called at x% of sites | % of genes called at x% of sites |
|---|---|---|
| 0% | 2,763 | 12.84 |
| 0-9% | 4,071 | 18.92 |
| >10% | 17,448 | 81.08 |
| >15% | 16,568 | 76.99 |
| 10-19% | 2,337 | 10.86 |
| >20% | 15,111 | 70.22 |
| >25% | 12,847 | 59.7 |
| 20-29% | 5,190 | 24.12 |
| >30% | 9,921 | 46.1 |
| >35% | 6,824 | 31.71 |
| 30-39% | 5,733 | 26.64 |
| >40% | 4,188 | 19.46 |
| >45% | 2,206 | 10.25 |
| 40-49% | 3,079 | 14.31 |
| >50% | 1,109 | 5.15 |
| >55% | 465 | 2.16 |
| 50-59% | 957 | 4.45 |
| >60% | 152 | 0.71 |
| >65% | 40 | 0.19 |
| 60-69% | 144 | 0.67 |
| >70% | 8 | 0.04 |
| >75% | 2 | 0.01 |
| 70-79% | 8 | 0.04 |
| >80% | 0 | 0 |
| >85% | 0 | 0 |
| 80-89% | 0 | 0 |
| >90% | 0 | 0 |
| >95% | 0 | 0 |
| 90-99% | 0 | 0 |
| 100% | 0 | 0 |

**2X no damage-masked**

| % of sites at 2X | # of genes called at x% of sites | % of genes called at x% of sites |
|---|---|---|
| 0% | 2,758 | 12.82 |
| 0-9% | 3,883 | 18.04 |
| >10% | 17,636 | 81.96 |
| >15% | 16,937 | 78.71 |
| 10-19% | 1,718 | 7.98 |
| >20% | 15,918 | 73.97 |
| >25% | 14,390 | 66.87 |
| 20-29% | 3,670 | 17.05 |
| >30% | 12,248 | 56.92 |
| >35% | 9,828 | 45.67 |
| 30-39% | 4,865 | 22.61 |
| >40% | 7,383 | 34.31 |
| >45% | 5,181 | 24.08 |
| 40-49% | 3,976 | 18.48 |
| >50% | 3,407 | 15.83 |
| >55% | 2,066 | 9.6 |
| 50-59% | 2,153 | 10.01 |
| >60% | 1,254 | 5.83 |
| >65% | 697 | 3.24 |
| 60-69% | 893 | 4.15 |
| >70% | 361 | 1.68 |
| >75% | 149 | 0.69 |
| 70-79% | 315 | 1.46 |
| >80% | 46 | 0.21 |
| >85% | 11 | 0.05 |
| 80-89% | 44 | 0.2 |
| >90% | 2 | 0.01 |
| >95% | 1 | 0.005 |
| 90-99% | 2 | 0.01 |
| 100% | 0 | 0 |
